# Live-imaging of endothelial Erk activity reveals dynamic and sequential signalling events during regenerative angiogenesis

**DOI:** 10.1101/2020.08.17.254912

**Authors:** Kazuhide S. Okuda, Mikaela Keyser, David B. Gurevich, Caterina Sturtzel, Scott Patterson, Huijun Chen, Mark Scott, Nicholas D. Condon, Paul Martin, Martin Distel, Benjamin M. Hogan

## Abstract

The formation of new blood vessel networks occurs via angiogenesis during development, tissue repair and disease. Angiogenesis is regulated by intracellular endothelial signalling pathways, induced downstream of Vascular endothelial growth factors (VEGFs) and their receptors (VEGFRs). A major challenge in understanding angiogenesis is interpreting how signalling events occur dynamically within endothelial cell populations during sprouting, proliferation and migration. Erk is a central downstream effector of Vegf-signalling and reports the signalling that drives angiogenesis. We generated a vascular Erk biosensor transgenic line in zebrafish using a kinase translocation reporter that allows live-imaging of Erk-signalling dynamics. We demonstrate the utility of this line to live-image Erk activity during physiologically relevant angiogenic events. Further, we reveal dynamic and sequential endothelial cell Erk-signalling events following blood vessel wounding. Initial signalling is dependent upon Ca^2+^ in the earliest responding endothelial cells, but is independent of Vegfr-signalling and local inflammation. The sustained regenerative response however, involves a Vegfr-dependent mechanism that initiates concomitant with the wound inflammatory response. This work thus reveals a highly dynamic sequence in regenerative angiogenesis that was not previously appreciated. Altogether, this study demonstrates the utility of a unique biosensor strain for analysing dynamic endothelial Erk-signalling events and validates a new resource for the study of vascular signalling in real-time.

## Introduction

The formation of new blood vessels from pre-existing vasculature (angiogenesis) is a fundamental process central in the formation of a viable embryo and in the pathogenesis of many diseases (Carmeliet and Jain, 2011;Chung and Ferrara, 2011;Potente et al., 2011). Angiogenesis is controlled by intricately regulated cell-cell, cell-matrix and intracellular signalling events. The activity of extracellular signal-regulated kinase (ERK) downstream of the vascular endothelial growth factor A (VEGFA)/VEGF receptor 2 (VEGFR2) signalling pathway is essential for both developmental and pathological angiogenesis (Koch and Claesson-Welsh, 2012;Simons et al., 2016). ERK-signalling is also required downstream of VEGFC/VEGFR3 signalling in lymphangiogenesis (Deng et al., 2013). ERK is required for angiogenic sprouting, proliferation and migration, with genetic or pharmacological inhibition of ERK-signalling resulting in impaired blood vessel development in zebrafish and mice (Srinivasan et al., 2009;Costa et al., 2016;Nagasawa-Masuda and Terai, 2016;Shin et al., 2016). Cancer-associated vessels have high ERK activity and inhibition of ERK-signalling blocks cancer-associated angiogenesis in mice (Wilhelm et al., 2004;Murphy et al., 2006). Beyond the formation of new vessels, ERK-signalling is also essential to maintain vascular integrity in quiescent endothelial cells (ECs) (Ricard et al., 2019), altogether demonstrating a central role for ERK in vascular biology.

Despite its importance, vascular ERK-signalling has largely been examined with biochemical assays or in tissues in static snapshots. Numerous studies have suggested that ERK-signalling is likely to be highly dynamic during angiogenic events, for example studies that examine Erk activation using antibodies to detect phosphorylated Erk (pErk) have observed changes associated with increased EC signalling, EC motility and specific EC behaviours (Costa et al., 2016;Nagasawa-Masuda and Terai, 2016;Shin et al., 2016). In zebrafish, live-imaging of blood ECs at single cell-resolution coupled with carefully staged immunofluorescence staining for pErk suggested that an underlying dynamic Erk-signalling event may control tip-cell maintenance in angiogenesis (Costa et al., 2016). Nevertheless, EC signalling dynamics at the level of key intracellular kinases, such as ERK, remain poorly understood. This gap in our understanding is largely due to a gap in our ability to live-image changes in signalling as they occur.

A number of new biosensors have now been applied *in vitro* and *in vivo* that allow live-imaging of proxy readouts for intracellular signalling events (reviewed in detail in (Shu, 2020)). One approach used, has involved application of biosensors that utilise fluorescence resonance energy transfer (FRET)-based readouts. The first ERK FRET-based biosensor (ERK activity reporter (EKAR)) was developed in 2008 (Harvey et al., 2008). Since then, modifications had been made to improve its sensitivity and dynamic range to generate other ERK FRET-based biosensors such as EKAR-EV, RAB-EKARev, and sREACh (Komatsu et al., 2011;Ding et al., 2015;Tang and Yasuda, 2017;Mehta et al., 2018). Importantly, these ERK FRET-based biosensors had been applied *in vivo* to visualise ERK-signalling dynamics in various cell types during development, cell migration, and wound healing (Kamioka et al., 2012;Mizuno et al., 2014;Goto et al., 2015;Hiratsuka et al., 2015;Kamioka et al., 2017;Takeda and Kiyokawa, 2017;Sano et al., 2018;Wong et al., 2018). While these ERK FRET-based biosensors have been widely reported, they are limited in requiring extensive FRET controls and a low speed of acquisition for FRET based imaging. More recently, Regot and colleagues generated the ERK-kinase translocation reporter (KTR)-Clover construct (hereafter referred to as EKC), that allows for dynamic analysis of ERK activity using a readout not involving FRET. A fluorescence-based kinase activity reporter translates ERK phosphorylation events into a nucleo-cytoplasmic shuttling event of a synthetic reporter (Regot et al., 2014). Thus, the KTR system allows rapid quantifiable measurements of ERK activity based upon subcellular localisation of a fluorescent fusion protein, and is more sensitive to phosphatase-mediated kinase activity downregulation when compared to other commonly used kinase activity reporters. This has been applied to enable dynamic ERK-signalling pulses to be analysed in single-cell resolution both *in vitro* and also *in vivo* (Regot et al., 2014;de la Cova et al., 2017;Mayr et al., 2018;Goglia et al., 2020), where it has demonstrated to be of high utility.

In this study, we generated a vascular EC-restricted EKC zebrafish transgenic strain and assessed its utility to study angiogenesis *in vivo*. We apply real-time quantification of Erk-signalling dynamics during developmental angiogenesis and vessel regeneration. We establish methods to quantify Erk activity during real time imaging that will be applicable in a variety of settings in vascular biology and beyond. Demonstrating the promise of this approach, we here identify an immediate early Erk-signalling response to wounding of vasculature that is Ca^2+^ signalling dependent and distinct from a later Vegfr-driven regenerative response. Overall, this work reports a unique resource for imaging of vascular signalling and further illuminates mechanisms of vascular regeneration following wounding.

## Results

### Generation of a zebrafish EC EKC transgenic line

KTRs utilise a kinase docking and target site that is juxtaposed to a phospho-inhibited nuclear localization signal (NLS) and attached to a fluorescent tag (Regot et al., 2014). Upon kinase activity the NLS is inactive and the fluorescent tag detected in the cytoplasm; when the kinase is not active dephosphorylated NLS leads to increased nuclear localisation. The EKC module that we took advantage of here relies upon the well characterised ERK-dependent transcription factor ELK1, utilising the ERK docking site (**Figure 1A**) (Chang et al., 2002;Regot et al., 2014). This reporter has previously been shown to report Erk activity *in vivo* (de la Cova et al., 2017;Mayr et al., 2018). To visualise real-time Erk-signalling in ECs, we expressed this reporter under the control of an EC-specific promoter (*fli1aep* (Villefranc et al., 2007)) (**Figures 1A-E**). Blood vessel development was unaffected in *Tg(fli1aep:EKC)* transgenic embryos and larvae (**Figures 1B-E**). Furthermore, transgenic adults displayed no adverse morphological features and were fertile (data not shown), indicating that EKC does not inhibit Erk-signalling *in vivo,* or cause developmental phenotypes and consistent with previous findings (Mayr et al., 2018).

**Figure 1:**
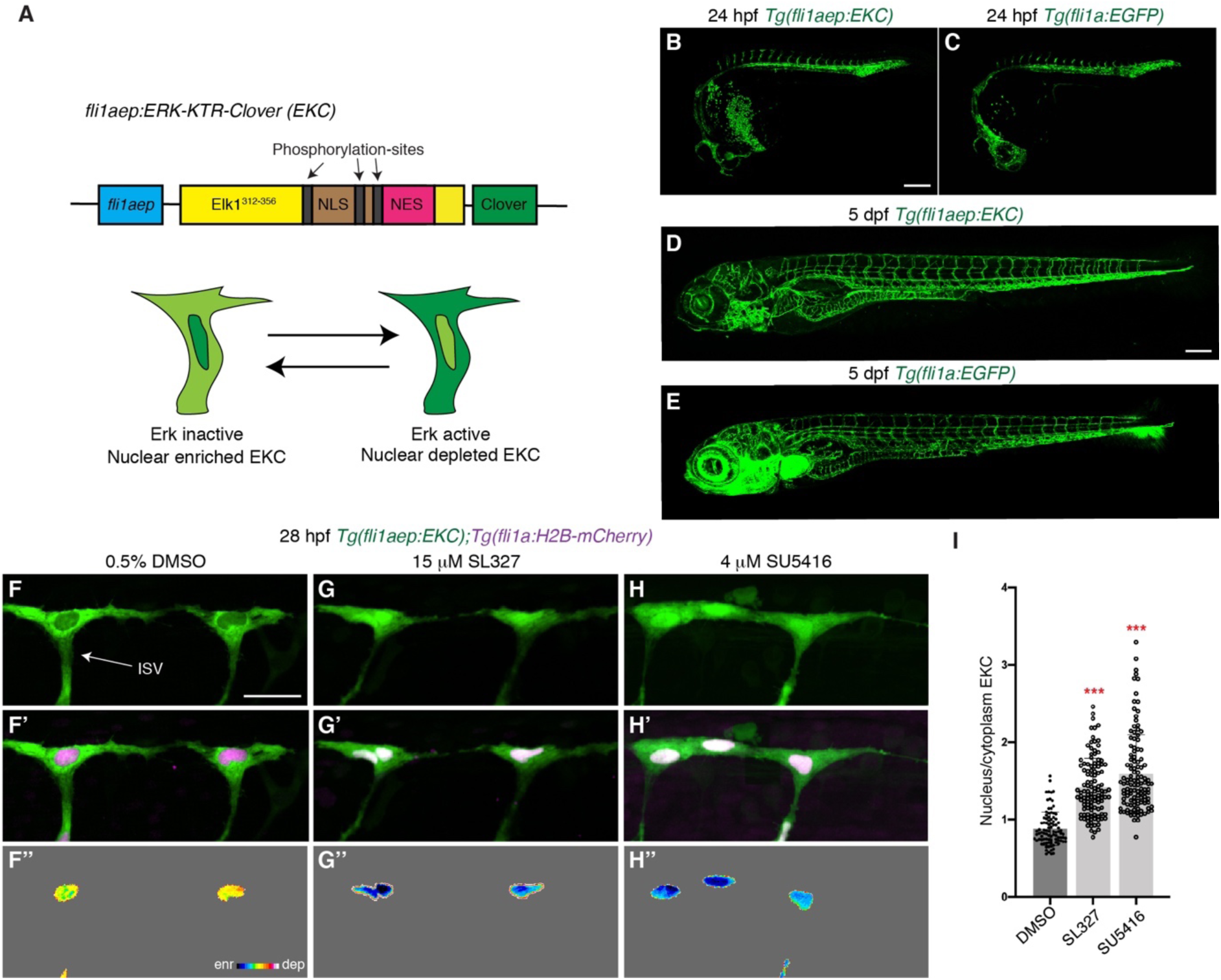
The *Tg(fli1aep:EKC)* transgenic line enables quantification of vascular Erk activity during development. (**A**) Schematic representation of the *fli1aep:ERK-KTR-Clover* (EKC) construct, and ECs with nuclear enriched EKC (bottom left, inactive Erk-signalling) and nuclear depleted EKC localisation (bottom right, active Erk-signalling). (**B-E**) Lateral confocal images of *the Tg(fli1aep:EKC)* (B,D) and *Tg(fli1a:EGFP)* (C,E) embryos/larvae at 24 hpf (B,C) and 5 dpf (D,E). (**F**-**H’’**) Lateral spinning disc confocal images of ISV ECs in 28 hpf *Tg(fli1aep:EKC);Tg(fli1a:H2B-mCherry)* embryos treated for 1 hour with either 0.5% DMSO (F-F’’), 15 μM SL327 (G-G’’), or 4 μM SU5416 (H-H’’). Images F-H show the *fli1aep:EKC* expression, while images F’-H’ show both the *fli1aep:EKC* and the *fli1a:H2B-mCherry* expression. Images F’’-H’’ show the nuclear *fli1aep:EKC* expression with intensity difference represented in 16 colour LUT (Fiji). The *fli1a:H2B-mCherry* signal was used to mask the nucleus. (**I**) Quantification of nucleus/cytoplasm EKC intensity in ISV tip ECs of 28 hpf embryos treated with either 0.5% DMSO (0.881, 93 ISV tip ECs, n=20 embryos), 15 μM SL327 (1.419, 114 ISV tip ECs, n=27 embryos), or 4 μM SU5416 (1.591, 118 ISV tip ECs, n=27 embryos). ISV: intersegmental vessel. Statistical test: Kruskal Wallis test was conducted for graph I. Error bars represent standard deviation. Scale bars: 200 μm for images B and D, 25 μm for image F.

To test if the *Tg(fli1aep:EKC)* line reports vascular Erk-signalling, embryos were treated with either DMSO, mitogen-activated protein kinase kinase (MEK) inhibitor SL327, or pan-VEGFR inhibitor SU5416, and vascular EKC localisation examined at 27 hpf. Tip ECs in developing ISVs had been shown to have high Erk activity (Costa et al., 2016;Nagasawa-Masuda and Terai, 2016;Shin et al., 2016) and we observed highly nuclear depleted EKC localisation in ISV tip cells in DMSO treated embryos. (**Figures 1F-F’’,I**). In contrast, ISV tip ECs of embryos treated with either SL327 or SU5416 had nuclear enriched EKC localisation indicating inactive Erk-signalling (**Figures 1G-I**). To best visualise these differences in signalling and differences shown below, we used a heat map of nuclear EC EKC intensity that is inverted so that blue-scale indicated low signalling (nuclear enriched) and red-scale indicates high signalling (nuclear depleted) (**Figures 1F’’-H’’**). Therefore, we confirmed that the *Tg(fli1ep:EKC)* (hereafter EC EKC) transgenic enables quantification of Erk activity in developing ECs.

### The EC EKC line enables visualisation and quantification of dynamic Erk activity during primary angiogenesis

We next sought to determine whether the EC EKC sub-cellular localisation reports physiologically relevant Erk-signalling events. Using immunofluorescence staining, ISV tip cells that sprout from the dorsal aorta (DA) have been shown to have higher Erk-signalling than ECs that remain in the DA during the initiation of angiogenesis (Nagasawa-Masuda and Terai, 2016;Shin et al., 2016). We examined 22 hpf embryos and indeed observed that sprouting ISV ECs display high Erk activity based upon EKC localisation (**Figure 1-figure supplement 1A-B**). However, many DA ECs also appeared to have nuclear depleted EKC localisation (**Figure 1-figure supplement 1A**, yellow arrows). To compare EKC and Erk-signalling levels between sprouting tip-cells and the DA, we utilised multiple methods. We found that measuring the nuclear/cytoplasm EKC intensity ratio in DA ECs was inaccurate because DA ECs are tightly packed, making cytoplasmic quantification unreliable (**Figure 1-figure supplement 1A’**). Previous studies have compared nuclear EKC with nuclear H2B-mCherry intensity in the same cell as a ratio to measure Erk activity (eg. in vulval precursor cells in the worm (de la Cova et al., 2017)). We assessed the ratio of nuclear EKC/H2B-mCherry intensity in double transgenic *Tg(fli1ae:EKC);Tg(fli1a:H2B:mCherry)* (hereafter referred to as EC-EKC/mCherry) embryos and found that the ISV tip cells had higher Erk activity than adjacent DA “stalk” ECs (**Figure 1-figure supplement 1A’’ and C**). We used a stable *Tg(fli1a:H2B-mCherry)* transgenic line with consistent H2B-mCherry intensity within ECs. Next, we investigated whether nuclear EKC intensity alone was sufficient to compare Erk-signalling between ECs. The ratio of nuclear EKC intensity of the sprouting ISV tip-cell and the adjacent DA “stalk” EC clearly showed higher signalling in tip-cells and was consistent with EKC/H2B-mCherry measurements (**Figure 1-figure supplement 1C**). Thus, we establish that both methods can be reliably used, when measurement of nuclear/cytoplasm EKC intensity is challenging because of the need to confidently identify a cells cytoplasm. We typically compare nuclear EKC intensities for subsequent analyses here.

Next, we correlated an ECs Erk-signalling state (based on EC EKC intensity) with its motility state (based on nuclear elongation) as previous studies have suggested a correlation (Costa et al., 2016). At 28 hpf, ISV tip cells were either located above the horizontal myoseptum with elongated nuclei indicative of high motility (migrating EC), or at the level of the future dorsal longitudinal anastomotic vessel, with rounded nuclei (non-migrating EC) (**Figure 1-supplement 1D-F**). We found that migrating ECs had higher Erk activity than non-migrating ECs, irrespective of their tip or stalk cell location in an ISV (**Figure 1-figure supplement 1D-H**). This is consistent with previous studies of Vegfa/Kdr/Kdrl/Erk signalling in zebrafish ISVs (Yokota et al., 2015;Shin et al., 2016) and confirms a strong correlation between ISV EC motility and EC Erk-signalling.

Using carefully staged immunofluorescence analyses, it was previously suggested that when tip cells divide in ISV angiogenesis, daughter cells show asymmetric Kdrl/Erk signalling that re-establishes the tip/stalk EC hierarchy (Costa et al., 2016). However, an analysis of fixed material can only ever imply underlying dynamics. To investigate the dynamics of Erk-signalling upon tip-cell division, we performed high-speed time-lapse imaging of ISV tip ECs as they undergo mitosis in 24 hpf embryos. Immediately preceding cell-division, ECs display cytoplasmic localisation of H2B- mCherry due to the disruption of the nuclear membrane (**Figure 2A**, yellow arrow). High-speed live-imaging of ISV tip ECs revealed nuclear enriched EKC localisation during cell division (**Figures 2A-C**), which was maintained until cytokinesis (**Figure 2B, Video 1**) but may reflect nuclear membrane breakdown rather than altered cellular signalling. Subsequently, daughter ECs in the tip position progressively increased their Erk activity, while ECs in the trailing stalk daughter position remained nuclear enriched, showing asymmetric Erk-signalling activity rapidly following cell division (**Figure 2B-I, Video 1**). To accurately assess this across multiple independent tip-cell divisions, we measured the ratio of tip/stalk daughter cell nuclear EKC intensity over time. This revealed that tip cells consistently increased their Erk activity relative to stalk cells in a progressive manner with the most dramatic asymmetry observed ∼21 minutes post-cytokinesis (**Figures 2B-K, Video 1**). Collectively, the EC EKC line enabled quantitative assessment of physiologically relevant Erk activity by real-time live imaging and confirmed previously suggested asymmetric signalling post tip-cell division.

**Figure 2:**
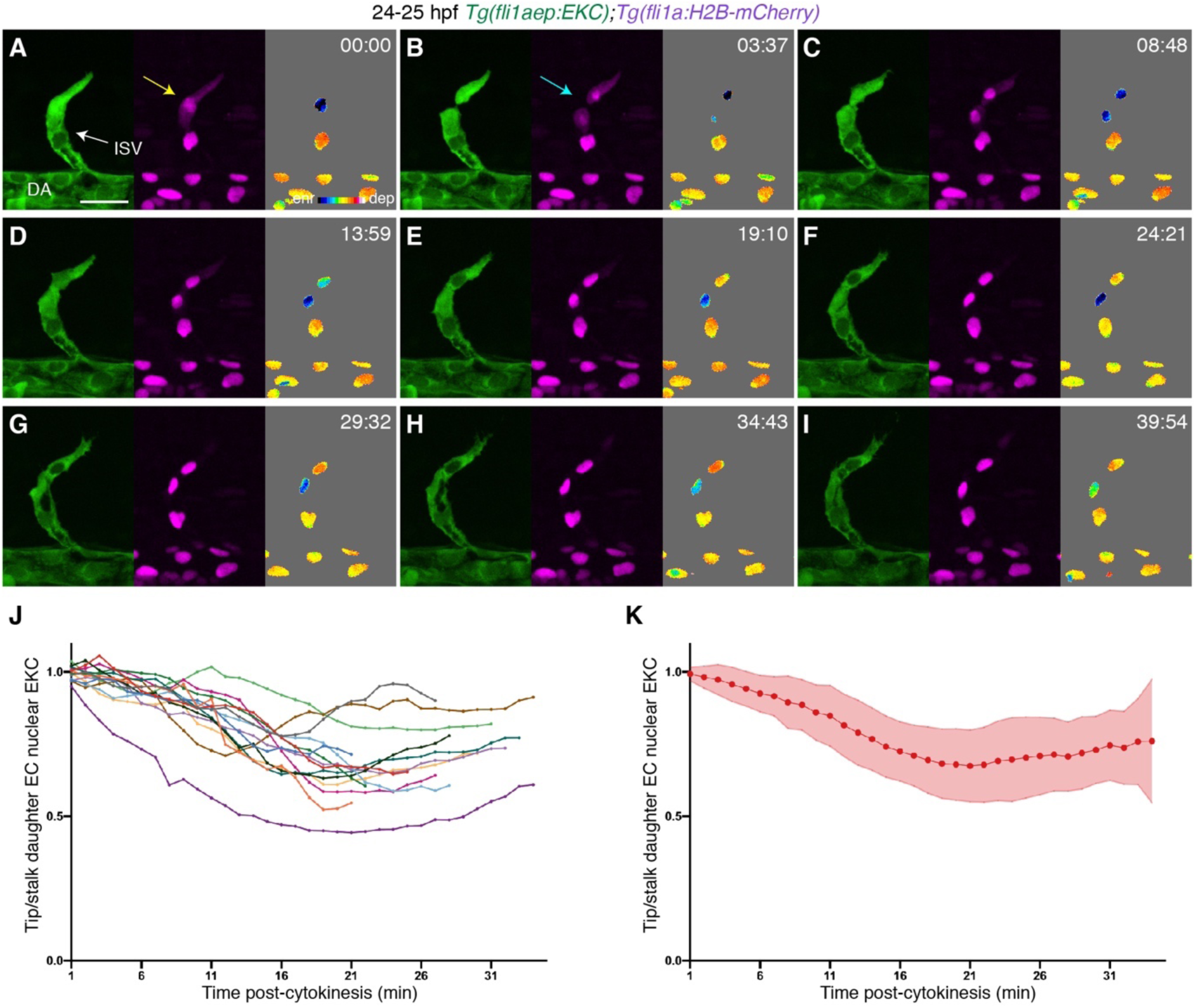
Tip cells show asymmetric Erk activity immediately following cell division. (**A-I**) Still images from Video 1 showing ISV ECs in a 24-25 hpf *Tg(fli1aep:EKC);Tg(fli1a:H2B-mCherry)* embryo at indicated time points. Left panels show *fli1aep:EKC* expression, middle panels show the *fli1a:H2B-mCherry* expression, and right panels show the nuclear *fli1aep:EKC* expression with intensity difference represented in 16 colour LUT (Fiji). The *fli1a:H2B-mCherry* signal was used to mask the nucleus. The yellow arrow indicates a tip ISV EC with cytoplasmic *fli1a:H2B-mCherry* expression. The light blue arrow indicates a tip ISV EC that has undergone cytokinesis. (**J,K**) Quantification of tip/stalk nuclear EKC intensity of daughter ECs post-cytokinesis (14 EC division events, n=14 embryos). Graph J shows quantification of individual biological replicates and graph K shows the average of all biological replicates. ISV: intersegmental vessel; DA: dorsal aorta. Error bars represent standard deviation. Scale bar: 25 μm.

### Vessel wounding induces rapid Erk activation

As a major downstream target for VEGFA/VEGFR2 signalling, ERK phosphorylation is essential for stimulating ectopic sprouting of otherwise quiescent mature vessels (Wilhelm et al., 2004;Murphy et al., 2006). However, Erk-signalling dynamics during pathological angiogenesis have not been analysed in detail. To determine whether the EC EKC line can be used to dynamically visualise Erk activation in ECs in pathological settings, we analysed EC Erk activity following vessel wounding using a laser ablation method. We chose this model because vessel wounding in 4 dpf larvae results in highly reproducible Vegfa/Kdr/Kdrl signalling-dependent vessel regeneration (Gurevich et al., 2018). Importantly, cell wounding induces rapid ERK-signalling waves *in vitro* and *in vivo* in other settings (Matsubayashi et al., 2004;Li et al., 2013;Hiratsuka et al., 2015;Aoki et al., 2017;Mayr et al., 2018).

To visualise Erk-signalling dynamics following cellular ablation and vessel wounding, we time-lapse imaged both ablated ISV ECs and the adjacent non-ablated ISV ECs in 4 dpf EC-EKC/mCherry larvae for 20 minutes before and for 22 minutes after vessel wounding. As a control, ISV ECs of unablated 4 dpf larvae were time-lapse imaged for the same period. EKC localisation in the majority of ISV ECs indicated low basal Erk-signalling in ECs of mature vessels (**Figures 3A,A’,C,C’,E,F, Videos 2-5**). Upon vessel wounding, Erk activity in ablated ISV ECs immediately increased (**Figures 3B,B’,E,F, Videos 3 and 4**). Surprisingly, Erk activity in ECs of ISVs located adjacent to the ablated ISV (termed adjacent ISV) also rapidly increased (**Figures 3D,D’,E,F, Videos 3 and 5**). Although the activation of Erk-signalling in adjacent ISV ECs was slightly slower than in ablated ISV ECs, the level of Erk activation in ablated and adjacent vessels was comparable by 15 minutes post-ablation (mpa, green dotted line) and consistent up to 22 mpa (**Figures 3F**). Both venous and arterial ECs equally showed Erk activation 15 mpa in ablated ISVs post-vessel wounding, suggesting that both venous and arterial ECs are able to rapidly activate Erk-signalling (**Figure 3G**).

**Figure 3:**
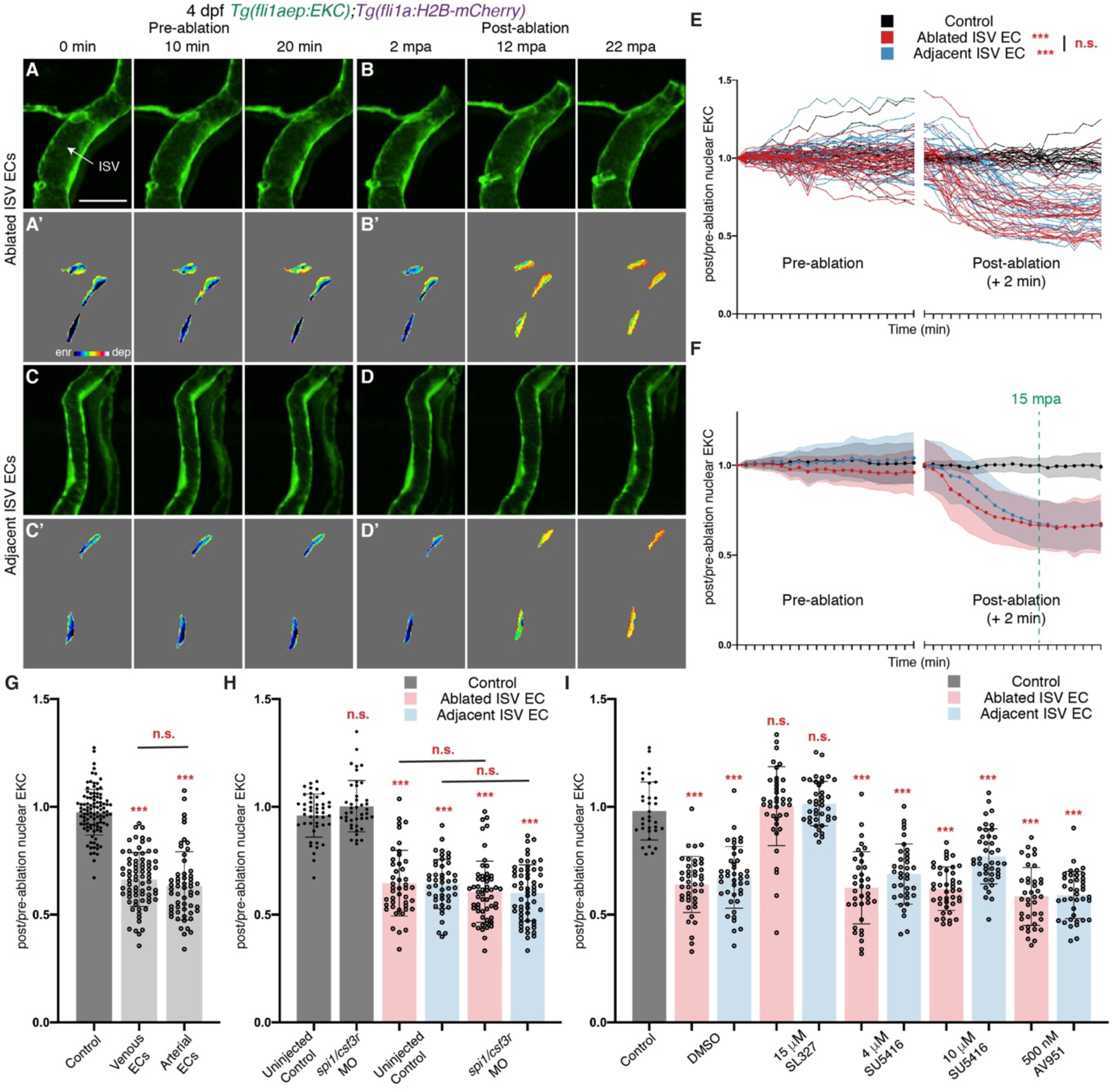
Wounded vessels rapidly activate Erk independent of macrophages of Vegfr-signalling. (**A-D’**) Still images from **Video 4** (A-B’) and **Video 5** (C-D’) showing ISV ECs of a 4 dpf *Tg(fli1aep:EKC);Tg(fli1a:H2B-mCherry)* larva at indicated time points before (pre-ablation) and after (post-ablation) vessel wounding. Erk activity is induced rapidly in wounded and unwounded, adjacent vessels. Images A-D show the *fli1aep:EKC* expression, while images A’-D’ show the nuclear *fli1aep:EKC* expression with intensity difference represented in 16 colour LUT (Fiji). The *fli1a:H2B-mCherry* signal was used to mask the nucleus. (**E,F)** Quantification of rapid Erk activation by the ratio of post/pre-ablation nuclear EKC intensity of ECs in non-ablated control ISVs (black, 24 ECs, n=8 larvae), ablated ISVs (red, 27 ECs, n=9 larvae), and adjacent ISVs (light blue, 27 ECs, n=9 larvae) before and after vessel wounding. Graph E shows quantification of individual ECs and graph F shows the average of all biological replicates. Green dotted line indicates 15 minutes post-ablation (mpa). (G) Quantification of post/pre-ablation nuclear EKC intensity 15 mpa in ECs of non-ablated control ISVs (103 ECs, n=34 larvae), ablated venous ISVs (75 ECs, n=25 larvae), and ablated arterial ISVs (57 ECs, n=19 larvae). (H) Quantification of post/pre-ablation nuclear EKC intensity 15 mpa in ECs of non-ablated uninjected control ISVs (45 ECs, n=15 larvae), non-ablated *spi1*/*csf3r* morphant ISVs (42 ECs, n=14 larvae), uninjected control ISVs (45 ablated/adjacent ISV ECs, n=15 larvae), and *spi1*/*csf3r* morphant ISVs (56 ablated ISV ECs and 57 adjacent ISV ECs, n=19 larvae). (I) Quantification of post/pre-ablation nuclear EKC intensity 15 mpa in ECs of 0.5% DMSO-treated non-ablated control ISVs (33 ECs, n=11 larvae), and ISVs of larvae treated with either 0.5% DMSO (42 ablated/adjacent ISV ECs, n=14 larvae), 15 μM SL327 (39 ablated/adjacent ISV ECs, n=13 larvae), 4 μM SU5416 (36 ablated/adjacent ISV ECs, n=12 larvae), 10 μM SU5416 (42 ablated/adjacent ISV ECs, n=14 larvae), or 500 nM AV951 (42 ablated/adjacent ISV ECs, n=14 larvae). ISV: intersegmental vessel. Statistical test: Permutation test was conducted for graph E. Kruskal Wallis test was conducted for graphs G-I. n.s. represents not significant. Error bars represent standard deviation. Scale bar: 20 μm

### The initial rapid Erk-signalling response is not induced by macrophages or Vegfr activity

Macrophages recruited to a wound site have been shown to provide a local source of Vegfa that stimulates vessel regeneration (Gurevich et al., 2018). Therefore, we investigated whether macrophages are required for rapid Erk activation in ISV ECs. As previously reported (Gurevich et al., 2018), macrophage recruitment to wounded site was minimal at 15 mpa, while robust macrophage recruitment was observed 3 hours post-ablation (hpa), suggesting that macrophages may not contribute to rapid Erk activation (**Figure 3-figure supplement 1A-D**). We depleted macrophages by knocking down Spi-1 proto-oncogene b (Spi1b) and Colony stimulating factor 3 receptor (Csf3r) using established morpholinos (Rhodes et al., 2005;Ellett et al., 2011;Pase et al., 2012) (**Figure 3-figure supplement 1E-G**). The rapid EC Erk activation post-wounding was unaffected upon macrophage depletion (**Figure 3H**, **Figure 3-figure supplement 1H-S’**). We next tested whether Vegfr-signalling is required for this rapid Erk activation. Erk activation in both ablated and adjacent ISV ECs 15 mpa was blocked in larvae treated with SL327, indicating that it requires upstream Mek activation (**Figure 3I**, **Figure 3-figure supplement 2D-M’**). However, treatment with two independent and validated VEGFR-inhibitors, SU5416 (**Figure 1H-I**) and AV951 (**Figure 3-figure supplement 2A-C**), did not impair the rapid Erk activation at 15 mpa (**Figure 3I, Figure 3-figure supplement 2O-Z’**). Therefore, Erk activation in both ablated and adjacent ISV ECs 15 mpa is induced independently of either macrophages or Vegfr-signalling, suggesting an initial response to vessel wounding that has not been previously examined.

### Following the initial rapid Erk activation, signalling is progressively restricted to regenerating vessels

Previous studies have shown that local wounding induces a rapid burst in ERK-signalling in surrounding cells (Matsubayashi et al., 2004;Li et al., 2013;Hiratsuka et al., 2015;Aoki et al., 2017;Mayr et al., 2018). To determine whether the initial Erk activation in ISV ECs post-vessel wounding was maintained, Erk activity was followed over a longer time-course until 3 hpa, when robust macrophage recruitment was observed (**Figure 3-figure supplement 1C,D**). Erk activity was again increased upon vessel wounding in both ablated and adjacent ISV ECs at 15 mpa (**Figure 4A-D, Figure 4-figure supplement 1A-I’**). Erk activity was maintained until 30 mpa in adjacent ISV ECs, but then gradually decreased and returned to non-ablated control levels by 1 hpa (**Figure 4B-D**). By contrast, high Erk activity was maintained for the duration in ablated ISV ECs (**Figure 4A,A’,C,D**). To test if this difference in Erk activity was influenced by long-term time-lapse imaging, Erk-signalling was analysed in ISV ECs of 3 hpa larvae. Similar to the time-course analysis, Erk activity in ablated ISV ECs was high at 3 hpa, while ECs in adjacent ISVs were at non-ablated control level (**Figure 4-figure supplement 1J-N**).

**Figure 4:**
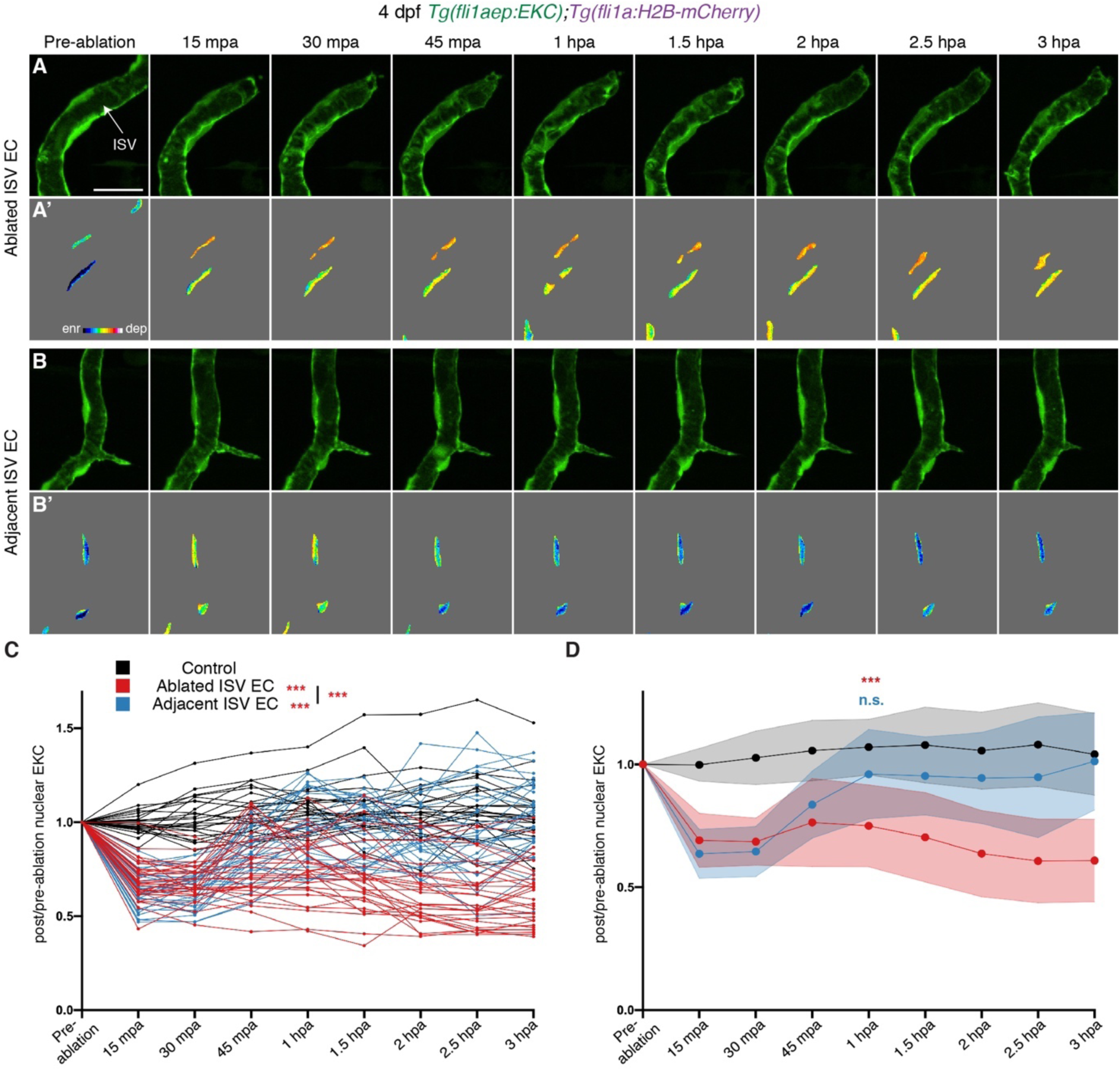
Wounded but not adjacent vessels retain high Erk activity as the regenerative response proceeds. (**A-B’**) Lateral spinning disc confocal images of ablated (A) and adjacent ISVs (B) of a 4 dpf *Tg(fli1aep:EKC);Tg(fli1a:H2B-mCherry)* larva before and following vessel wounding at indicated timepoints. Erk activity is progressively lost in the adjacent but retained in the wounded vessel. Images A and B show the *fli1aep:EKC* expression, while images A’ and B’ show the nuclear *fli1aep:EKC* expression with intensity difference represented in 16 colour LUT (Fiji). The *fli1a:H2B-mCherry* signal was used to mask the nucleus. (**C,D**) Quantification of post/pre-ablation nuclear EKC intensity of ECs in non-ablated control ISVs (black, 24 ECs, n=8 larvae), ablated ISVs (red, 30 ECs, n=10 larvae), and adjacent ISVs (light blue, 30 ECs, n=10 larvae) before and after vessel wounding at indicated timepoints. Graph C shows the quantification of individual ECs and graph D shows the average of all biological replicates. ISV: intersegmental vessel. Statistical test: Permutation test was conducted for graph C, Kruskal Wallis test was conducted for graph D. n.s. represents not significant. Error bars represent standard deviation. Scale bar: 20 μm

Given that the initial rapid burst of Erk activation progressively returns to basal levels in unwounded vessels, we assessed if this was a general wound response. We examined the initial Erk-signalling burst in muscle and skin cells following a large puncture wound using a ubiquitous EKC strain (Mayr et al., 2018). This confirmed that an initial activation of Erk signalling in cells surrounding the puncture wound was only transient (**Video 6**) and in this case was progressively lost even in cells at the immediate site of the wound, unlike in regenerating vessels. To further investigate whether only regenerating ISVs maintain high Erk activity after wounding, tissue in between the ISVs was ablated without injuring the ISVs in 4 dpf EC-EKC/mCherry larvae. Erk activity in surrounding ISV ECs was analysed at 15 mpa and 3 hpa. Similar to vessel ablation, this adjacent tissue ablation resulted in rapid activation of Erk-signalling in ISV ECs (**Figure 4-figure supplement 2A-C**). Erk activity in these ECs decreased to non-ablated control levels by 3 hpa (**Figure 4-figure supplement 2A-C**). Therefore, Erk-signalling is immediately activated in muscle, skin epithelial and ECs upon injury, but only regenerating vessels retain this high activity at 3 hpa.

### Vegfr-signalling drives ongoing Erk activity to control vessel regeneration

We next examined if ongoing Erk activity in ablated ISV ECs was maintained by Vegfr-signalling consistent with earlier reports (Gurevich et al., 2018). To test this, we analysed Erk activity of ablated ISV ECs in 3 hpa larvae treated with inhibitors of the Kdr/Kdrl/Mek/Erk signalling pathway. Treatment with SL327 inhibited Erk activation at 3 hpa, as did treatment with the Vegfr-inhibitor SU5416, indicating that Vegfr/Mek signalling is required for sustained high Erk activity in ablated ISV ECs 3 hpa (**Figure 5A, Figure 5-figure supplement 1A-F’,I-J’**). To determine the functional relevance of this in ongoing regeneration, we treated embryos following cell ablation-based vessel wounding with SU5416 or two independent Mek inhibitors: SL327 and Trametinib. We observed that inhibition of Vegfr- or Erk-signalling blocked all ongoing vessel regeneration (**Figure 5B-F**).

**Figure 5:**
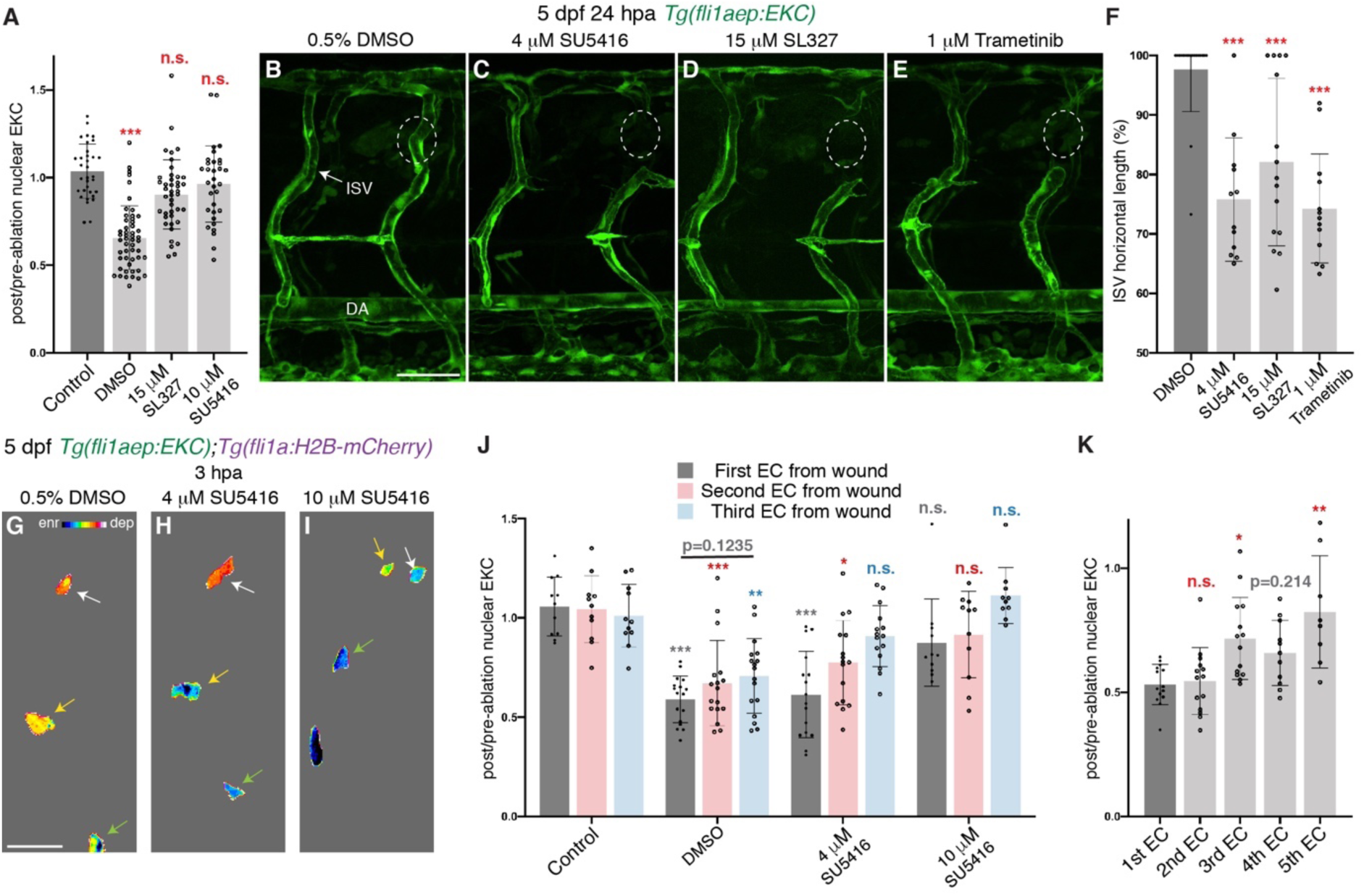
Erk activity in ablated vessels is maintained through the Vegfr pathway. (**A**) Quantification of post/pre-ablation nuclear EKC intensity 3 hpa in ECs of 0.5% DMSO-treated non-ablated control ISVs (33 ECs, n=11 larvae), and ablated ISVs of larvae treated with either 0.5% DMSO (51 ECs, n=17 larvae), 15 μM SL327 (42 ECs, n=14 larvae), 4 μM SU5416 (47 ECs, n=16 larvae), or 10 μM SU5416 (32 ECs, n=11 larvae). Ongoing signalling requires Vegfr and Erk activity. (**B-E**) Lateral spinning disc confocal images of 24 hpa 5 dpf *Tg(fli1ep:EKC)* larvae treated with either: 0.5% DMSO (B), showing a regenerated ISV; or 4 μM SU5416 (C), 15 μM SL327 (D), or 1 μM Trametinib (E); all of which blocked ISV regeneration. White circles show the wounded site of each larvae. (**F**) Quantification of ISV horizontal length percentage of ablated ISV in 24 hpa 5 dpf *Tg(fli1ep:EKC)* larvae treated with either 0.5% DMSO (n=18 larvae), 4 μM SU5416 (n=12 larvae), 15 μM SL327 (n=15 larvae), or 1 μM Trametinib (n=13 larvae). (**G-I**) Lateral spinning disc confocal images of ablated ISV ECs in 4 dpf 3 hpa *Tg(fli1aep:EKC);Tg(fli1a:H2B-mCherry)* larvae treated with either 0.5% DMSO (G), 4 μM SU5416 (H), or 10 μM SU5416 (I). Images show that the nuclear *fli1aep:EKC* expression was consistently higher and more Vegfr-dependent closer to the wound. Intensity difference represented in 16 colour LUT (Fiji). The *fli1a:H2B-mCherry* signal was used to mask the nucleus. Arrows indicate first (white), second (yellow), and third (green) ECs from the wounded site. Full images: **Figure 5-figure supplementary 1D’,H’,J’**. (J) Quantification of post/pre-ablation nuclear EKC intensity 3 hpa in first (dark grey), second (red) and third (light blue) ECs from the wounded site of 0.5% DMSO-treated non-ablated control ISVs (11 first, second and third ECs, n=11 larvae), and ablated ISVs of larvae treated with either 0.5% DMSO (17 first, second and third ECs, n=17 larvae), 4 μM SU5416 (16 first and second ECs, and 15 third ECs, n=16 larvae), or 10 μM SU5416 (11 first and second ECs, and 10 third ECs, n=11 larvae). Data were taken from graph A. (K) Quantification of post/pre-ablation nuclear EKC intensity 3 hpa in first (14 ECs, n=14 larvae), second (14 ECs, n=14 larvae), third (14 ECs, n=14 larvae), forth (11 ECs, n=11 larvae), and fifth (8 ECs, n=8 larvae) ECs from the wounded site of ablated ISVs in 4 dpf *Tg(fli1aep:EKC);Tg(fli1a:H2B-mCherry)* larvae. Data for the first, second, and third ECs were taken from **Figure 4-figure supplement 1N**. ISV: intersegmental vessel. DA: dorsal aorta. Statistical test: Kruskal Wallis test was conducted for graphs A,F,J,K. n.s. represents not significant. Error bars represent standard deviation. Scale bar: 50 μm for image B, 15 μm for image G.

Interestingly, we noted that while treatment with SU5416 at 10 μM blocked ongoing Erk activation (**Figure 5-figure supplement 1I-J’**), treatment with the same inhibitor at a lower dose of 4 μM did not completely block Erk activity (**Figure 5-figure supplement 1G-H’**). To further investigate this with more spatial resolution, we examined Erk activity in ISV ECs relative to their distance from the cellular ablation site. Erk-signalling in the first, second, and third ISV ECs from the wound was activated 3 hpa in control larvae, while treatment with 10 μM SU5416 inhibited signalling in ECs located in all of these positions (**Figure 5G,I,J, Figure 5-figure supplement 1C-D’,I-J’**). However, with the intermediate dose of 4 μM SU5416, while the closest cell to the wound site still displayed Erk activity, as did the second cell from the wound site, the third and furthest from the wounded sites were now inhibited (**Figure 5H,J, Figure 5-figure supplement 1G-H’**). These results suggest that there is a gradient of Vegfr/Erk signalling activity in the ablated ISV ECs resulting in higher Vegfr/Erk activity in ECs closer to the wounded site, which can only be inhibited with SU5416 at higher concentrations. To test this, we examined the EC EKC levels relative to cell position and directly confirmed this graded activation at 3 hpa (**Figure 5K, Figure 4-figure supplement 1J-M’**). Together, these analyses confirm that during the ongoing response to vessel wounding, Vegfr-signalling is crucial and drives a graded signalling response to regulate regenerating vessels.

### Ca^2+^ signalling is required for initial rapid Erk activation upon vessel wounding

Although Vegfr-signalling is required for sustaining high Erk activity in ablated ISV ECs, it is not required for inducing the initial rapid Erk-signalling response. Activated by ATP released by damaged cells, Ca^2+^ signalling is one of the first intra-cellular mechanisms to be activated post-wounding in many cell types (reviewed in detail in (Ghilardi et al., 2020)). Accordingly, mechanical injury of blood vessels has been shown *in situ* to rapidly activate Ca^2+^ signalling in neighbouring endothelial cells in excised rat aorta (Berra-Romani et al., 2008;Berra-Romani et al., 2012). Although Ca^2+^ signalling activates Erk-signalling in endothelial cells downstream of the Vegfa/Vegfr2 signalling pathway (Koch and Claesson-Welsh, 2012;Moccia et al., 2012), Ca^2+^ signalling alone can also activate Erk-signalling (Xiao et al., 2011;Handly et al., 2015).

To determine whether Ca^2+^ signalling is rapidly activated in ablated ISV ECs in our model, we measured the dynamic expression of a ubiquitously expressed GCamp, a GFP-based Ca^2+^ probe, using the *Tg(actb2:GCaMP6f);Tg(kdrl:mCherry-CAAX)* transgenic line (Herzog et al., 2019). ISVs in non-ablated 4 dpf larvae did not show Ca^2+^ signalling, indicating low Ca^2+^ activity in stable ISVs (**Figure 6B, Video 7**). In contrast, ablated ISV ECs showed a rapid pulse of active Ca^2+^ signalling at 5mpa, which progressively decreased and returned to the level of surrounding tissue (**Figure 6A,B, Video 8**). Similar to ISVs in non-ablated controls, active Ca^2+^ signalling was not observed in adjacent ISVs (**Figure 6A,B, Video 8**). To determine whether Ca^2+^ signalling is required for rapid Erk activation in ablated ISV ECs, 4 dpf EC- EKC/mCherry larvae were treated with either DMSO or a potent Ca^2+^ signalling inhibitor Nifedipine for 30 minutes. Nifedipine treatment did not inhibit Erk-signalling activation in adjacent ISV ECs resulting in similar Erk activity as DMSO treated larvae 15 mpa (**Figure 6C, Figure 6-figure supplement 1A-B’,G-J’**). However, Erk activation in ablated ISV ECs (where we observed the GCaMP signal above) was significantly reduced when compared to DMSO treated larvae (**Figure 6C, Figure 6-figure supplement 1C-F’**). This was reproduced in an independent experiment using Amlodipine, an alternative Ca^2+^ signalling inhibitor (**Figure 6D, Figure 6-figure supplement 1K-T’**). This indicates that Ca^2+^ signalling plays a crucial role upstream of Erk in the wound response, but also that the response is differentially regulated in ablated compared with adjacent vessels, indicative of additional underlying signalling complexity.

**Figure 6:**
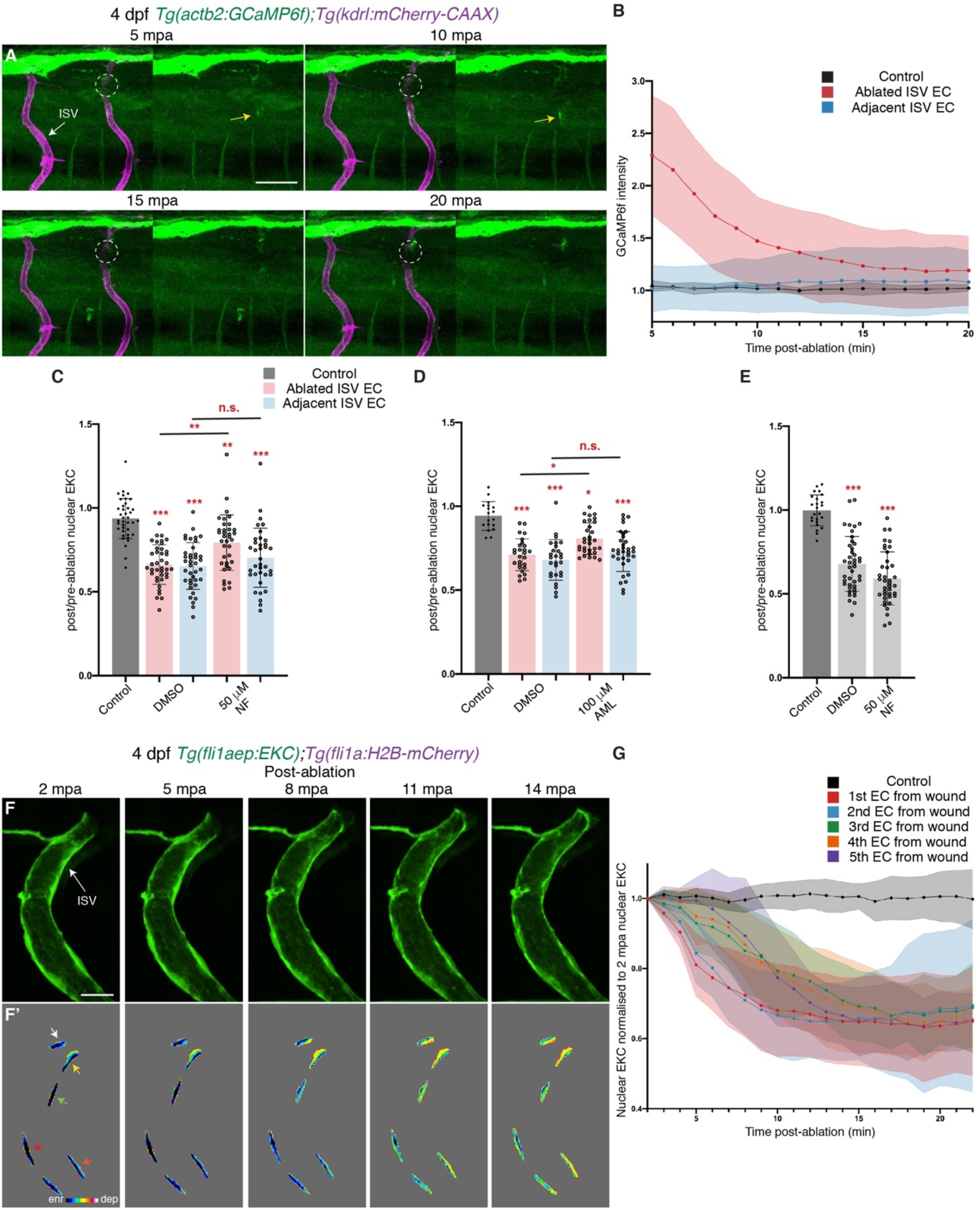
Ca^2+^ signalling is required for rapid Erk-activation in ablated vessels. (A) Still images from **Video 8** following vessel wounding and demonstrating a pulse of activation Ca2+ signalling immediately adjacent to the wound, at the indicated time points (4 dpf). Left panels show both the *actb2:GCaMP6f* and *kdrl:mCherry-CAAX* expression, while right panels show only the *actb2:GCaMP6f* expression. White circles show the wounded site. Yellow arrows show ISV ECs with active Ca^2+^ signalling. (B) Quantification of *actb2:GCaMP6f* intensity in unablated control ISVs (black, n=4 larvae), and ablated (red, n=10 larvae) and adjacent (light blue, n =10 larvae) ISVs following vessel wounding at indicated time points. Intensity was normalised to *actb2:GCaMP6f* intensity in unablated tissue in the same larvae. (C) Quantification of post/pre-ablation nuclear EKC intensity 15 mpa in ECs of 1% DMSO-treated non-ablated control ISVs (39 ECs, n=13 larvae), and ISVs of larvae treated with either 1% DMSO (39 ablated/adjacent ISV ECs, n=13 larvae) or 50 μM Nifedipine (36 ablated/adjacent ISV ECs, n=12 larvae). (D) Quantification of post/pre-ablation nuclear EKC intensity 15 mpa in ECs of 1% DMSO-treated non-ablated control ISVs (18 ECs, n=6 larvae), and ISVs of larvae treated with either 1% DMSO (27 ablated/adjacent ISV ECs, n=9 larvae) or 100 μM Amplopidine (31 ablated ISV ECs and 33 adjacent ISV ECs, n=11 larvae). (E) Quantification of post/pre-ablation nuclear EKC intensity 3 hpa in ECs of 1% DMSO-treated non-ablated control ISVs (24 ECs, n=8 larvae), and ablated ISVs of larvae treated with either 1% DMSO (42 ECs, n=14 larvae) or 50 μM Nifedipine (39 ECs, n=13 larvae). (**F,F’**) Still images from **Video 3** showing ablated ISV ECs of a 4 dpf *Tg(fli1aep:EKC);Tg(fli1a:H2B-mCherry)* larva at indicated time points after vessel wounding. Rapid activation of Erk is progressive from the wound site to the vessel base. Image F show the *fli1aep:EKC* expression, while image F’ shows the nuclear *fli1aep:EKC* expression at indicated timepoints with intensity difference represented in 16 colour LUT (Fiji). The *fli1a:H2B-mCherry* signal was used to mask the nucleus. Arrows indicate first (white), second (yellow), third (green), forth (red), and fifth (orange) ECs from the wounded site. (**G**) Quantification of nuclear EKC intensity normalised to the nuclear EKC intensity at 2 mpa in ECs of ISVs in non-ablated control larvae (black, 24 ECs, n=8 larvae), and the first (red, 9 ECs, n=9 larvae), second (blue, 9 ECs, n=9 larvae), third (green, 9 ECs, n=9 larvae), fourth (orange, 8 ECs, n=8 larvae), and fifth (purple, 5 ECs, n=5 larvae) ablated ISV ECs from the wounded site following vessel wounding. ISV: intersegmental vessel. Statistical test: Kruskal Wallis test was conducted for graphs C-E. n.s. represents not significant. Error bars represent standard deviation. Scale bars: 50 μm for image A, 15 μm for image F.

We next tested whether Ca^2+^ signalling is required for maintaining Erk activity in ablated ISV ECs 3 hpa. To assess ongoing signalling, 4 dpf EC-EKC/mCherry larvae were treated with either DMSO or Nifedipine 30 minutes prior to the 3 hpa timepoint. Activation of Erk-signalling in ablated ISV ECs 3 hpa was not inhibited by Nifedipine (**Figure 6E, Figure 6-figure supplement 1U-Z’**). Thus, Ca^2+^ signalling is required for rapid Erk activation, but not for maintaining Erk activity in ablated ISV ECs. In the analysis of Ca^2+^ signalling following vessel wounding, we noted that this transient pulse of Ca^2+^ signalling was highest in the ECs closest to the wounded site (**Video 8**). Thus, we further sought to determine if Erk-signalling in ECs closest to the wound activates first during the initial dynamic induction. Detailed analysis (**Video 3**), showed that Erk-signalling in ECs closer to the wounded site (first and second positioned ECs) activated first, followed by ECs further away from the wounded site (third, fourth and fifth ECs) (**Figure 6F-G**). This shows that like the initial burst in Ca^2+^ signalling post-vessel wounding, Erk-signalling is activated progressively in ECs closest to the wounded site first, followed by those further away.

## Discussion

ERK-signalling is a downstream target for a number of pathways essential for development (including VEGFA/VEGFR2, EGF/EGFR, FGF/FGFR pathways) and plays a central role in organ development by promoting proliferation, growth, migration and differentiation (Hogan and Schulte-Merker, 2017;Lavoie et al., 2020). As such, Erk-signalling must be tightly regulated in both its spatial and temporal activation. To understand how dynamically Erk activity is regulated in developing vasculature, we generated the *Tg(fli1aep:EKC)* transgenic line and validated its use as a proxy readout of active Erk-signalling in vasculature. We found that it both provided a valid readout for physiological Erk-signalling and uncovered previously unappreciated Erk-signalling dynamics during vessel regeneration (**Figure 7**). In the context of tip cell proliferation in angiogenesis, we revealed very rapid post-cell division signalling asymmetry, confirming previous work based on static imaging (Costa et al., 2016). In regenerative angiogenesis, we reveal a two-step mechanism for Erk-signalling activation post-vessel wounding, that involves an immediate and an ongoing signalling response that progressively limits Erk-signalling to vessels that are regenerating. Importantly, this study shows the utility of this new transgenic line to elucidate dynamic Erk-signalling events in vertebrate ECs and we suggest it will be a useful tool for diverse future studies of development and disease.

**Figure 7:**
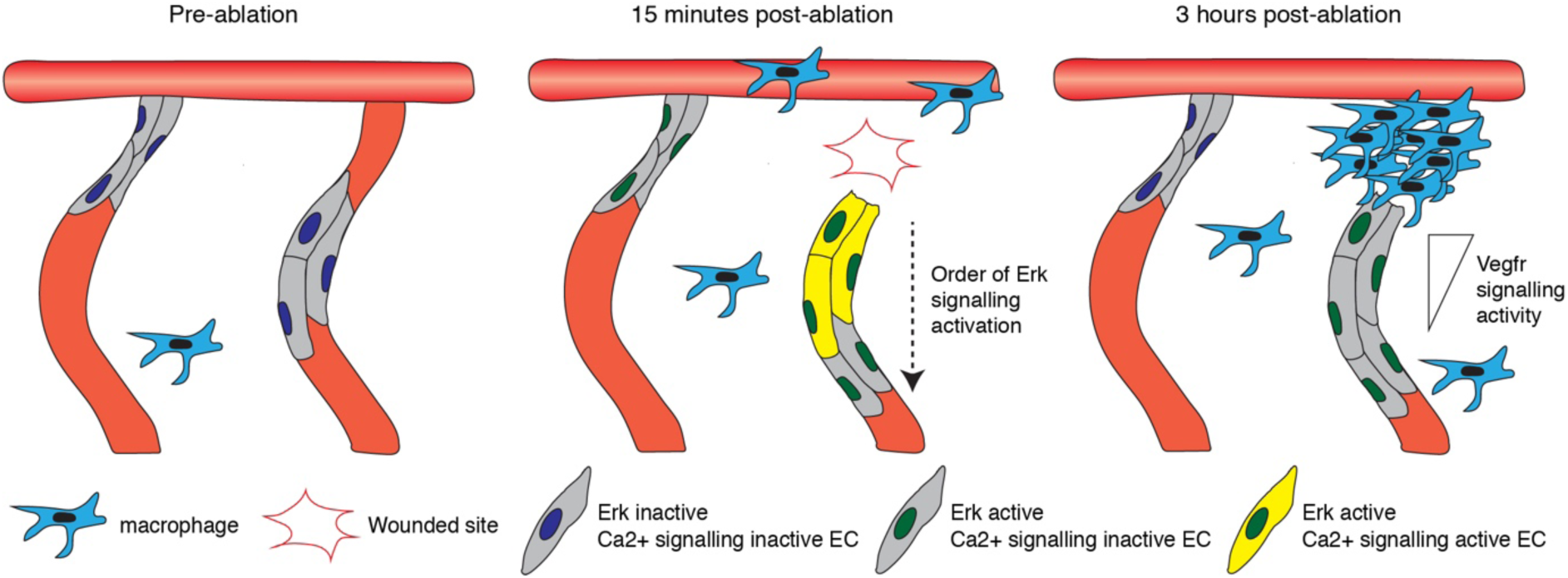
A two-step mechanism for activating and maintaining Erk activity in regenerating vessels. Schematic representation of the two-step mechanism employed by ECs to activate Erk-signalling following vessel wounding. Pre-ablation (left), the majority of ECs are Erk-signalling inactive. Following vessel wounding (middle), both ablated and adjacent ISV ECs rapidly activate Erk-signalling. Ca^2+^ signalling is also rapidly activated following vessel wounding but only in ablated ISV ECs, particularly in ECs close to the wounded site. Ca^2+^ signalling activity contributes to the activation of Erk-signalling in ablated ISV ECs in a sequential manner, starting from ECs close to the wounded site. Erk-signalling in adjacent ISV ECs has returned to pre-wound levels by 3 hpa (right). Erk activity in ablated ISV ECs is sustained through Vegfr-signalling, likely stimulated by Vegfa secreted from macrophages recruited to the wounded site. ECs closer to the wounded site are less sensitive to Vegfr signalling inhibition, suggesting higher signalling, when compared to ECs further away.

At a technical level, we used various quantification methods for measuring Erk activity in ECs and all generated valid results. The ratio of nuclear/cytoplasm EKC localisation gives the most accurate readout (Regot et al., 2014), but can only be used when cells are not tightly packed and their cytoplasmic fluorescence can be accurately measured. This is especially challenging for ECs which overlap and have unpredictable morphology in functional tubes. De la Cova and colleagues, recently applied ERK-nKTR, a second generation ERK KTR which includes a nuclear localised H2B-mCherry allowing them to quantify Erk activity based on Clover/mCherry in nuclei in *C. elegans* (de la Cova et al., 2017). We used a similar approach here with two independent transgenes driving EKC and H2B-mCherry and produced highly consistent results. It is worth noting that inter-embryo/larvae variations in H2B-mCherry intensity need to be considered when using this approach. Finally, we also used the measurement of nuclear EKC normalised to the average EKC intensity of the DA to normalise for embryo to embryo variation. This approach also provided data consistent with the other two methods. Thus, overall this EC EKC model is highly robust with multiple methods to quantify and normalise sensor localisation. As KTR reporters are used more frequently *in vivo* in the future, the quantification methods used here may be applied to many scenarios analysing cellular Erk activity in cells with complex 3D morphology.

Studies in zebrafish and xenopus have demonstrated rapid Erk activation in epithelial cells upon local wounding, which subsides relatively quickly (within 1hpa) as tissue repair progresses (Li et al., 2013;Mayr et al., 2018). Interestingly, our work shows a similar, very rapid, Erk activation in all vasculature in proximity of a wound. This suggests a common, initial, rapid Erk-signalling response immediately post-wounding in many different cell types and tissues – as if cells adjacent to a wound are rapidly primed to respond. However, in the vasculature this signalling returned to pre-ablation levels by 1 hpa, while Erk activity was maintained for a longer timeframe only in the wounded vessels. This ongoing, later signalling was maintained through Vegfr activity, likely stimulated by Vegfa secreted locally by recruited macrophages (Gurevich et al., 2018). Thus, Erk-signalling dynamics between wounded (ablated) and unwounded (adjacent) vessels differed significantly. We suggest this difference represents an initial priming of the wounded tissue (the rapid Erk response) that is replaced overtime with sustained vascular Erk-signalling that is essential in the regenerative response.

Rapid Ca^2+^ signalling post-wounding is observed in multiple systems *in vitro* and *in vivo* (reviewed in detail in (Ghilardi et al., 2020)). Using both quantitative live imaging and pharmacological inhibition, we found that Ca^2+^ signalling is required for Erk activation in ablated ISV ECs. Taking advantage of the high spatial and temporal resolution in our model, we found that Ca^2+^-dependent Erk-signalling is activated progressively from cells closest to the wound to cells further away. This may be consistent with a wave of tissue Ca^2+^ signalling through the wounded vessel. Activation of Erk-signalling at 2 mpa in wounded epithelial cells in xenopus promotes actomyosin contraction and wound closure (Li et al., 2013). Therefore, rapid Ca^2+^ signalling-mediated Erk activation in the wounded vessel may ensure efficient wound closure in ablated ISVs. However, we found no evidence that Ca^2+^ signalling influenced the broader, rapid Erk-signalling response in unwounded but adjacent vasculature. One interesting candidate to contribute to this broader mechanism is altered tissue tension associated with the tissue ablation, which had been shown in some contexts to modulate ERK-signalling (Rosenfeldt and Grinnell, 2000;Hirata et al., 2015). Perhaps consistent with this idea, we did not identify a mechanism required for rapid Erk activation in adjacent ISV ECs and vessel wounding was not required - tissue wounding in between ISVs alone activated Erk-signalling in surrounding ECs. Further work is needed to fully appreciate the mechanical tissue contributions in this response. Nevertheless, rapid Erk activation in ECs upon wounding seems likely to potentiate these ECs to more rapidly respond to external growth factors such as Vegfa upon the later activation of the inflammatory response and initiation of a sustained regenerative angiogenesis.

Taking advantage of spatial information in the imaging data, we showed that ECs in wounded ISVs that are actively regenerating at 3 hpa display a graded signalling response along the vessel at the level of Vegfr/Erk activity. This is likely due to a discrete local source of Vegfa from macrophages and may explain why unwounded ISV ECs, which are further away from the Vegfa source, do not sustain high Erk activity at 3 hpa. In bigger wounds, excessive angiogenesis has been previously reported to occur from adjacent ISVs and macrophage-dependent vascular regression is then required to ensure vessel patterns returns to their original state (Gurevich et al., 2018). Therefore, we hypothesise that maintaining Erk activity only in ECs of vessels that need to regenerate in this ablation model, ensures EC proliferation and migration only occurs in regenerating vessels, and prevents excessive angiogenesis. Further studies could investigate Erk-signalling dynamics of ECs in bigger wounds, which more closely resemble traumatic injury in humans and could further assess Erk-signalling dynamics in excessive angiogenesis and regression.

Blood vessels constantly remodel to accommodate for the needs of the human body during development and disease (Carmeliet and Jain, 2011;Chung and Ferrara, 2011;Potente et al., 2011). It is therefore not surprising for Erk-signalling, which is a key modulator of angiogenesis, to be highly dynamic in ECs. As a novel tool that allows real-time analysis of Erk activity, EC EKC biosensors will be useful for elucidating Erk-signalling events in vasculature in an array of settings and different vertebrate models. Importantly, in zebrafish the *Tg(fli1aep:EKC)* transgenic line can be coupled with both established and novel mutants with vascular phenotypes to investigate how real-time EC Erk-signalling dynamics is affected in the absence of key vascular genes. Further, dynamic Erk-signalling events in ECs in zebrafish disease models associated with increased angiogenesis such as in cancer (Nicoli et al., 2007) and tuberculosis (Oehlers et al., 2015) can be analysed using this EC EKC model. This could highlight novel pathological Erk-signalling events in ECs, that could be normalised using drugs shown to modulate Erk-signalling (Goglia et al., 2020). Of note, KTR constructs for other kinases such as AKT, JNK and p38 are also now available (Regot et al., 2014;Maryu et al., 2016). Also, other types of fluorescence-based kinase activity reporters such as separation of phases-based activity reporter of kinases (SPARK), that had been shown to work in zebrafish could be applied to visualise EC Erk activity in zebrafish (Zhang et al., 2018). Future studies should combine multiple zebrafish transgenics expressing such reporters to elucidate real-time interaction between different kinases in signalling pathways and to understand the real-time mechanistic drivers of development and disease.

## Materials and methods

### Key resources table

| **Reagent type**<br>**(species) or**<br>**resource** | **Designation** | **Source or**<br>**reference** | **Identifiers** | **Additional information/reagent source** |
| --- | --- | --- | --- | --- |
| Genetic reagent<br>(*D. rerio*) | *Tg(fli1a:H2B-*<br>*mCherry)<sup>uq37bh</sup>* | (Baek et al.,<br>2019) | ZFIN ID: ZDB-ALT-191011-5 | Ben M Hogan (Organogenesis and Cancer<br>Program, Peter MacCallum Cancer Centre,<br>Australia) |
| Genetic reagent<br>(*D. rerio*) | *Tg(fli1a:EGFP)<sup>y1</sup>* | (Lawson and<br>Weinstein, 2002) | ZFIN ID: ZDB-ALT-011017-8 | Brant M Weinstein (National Institute of Child<br>Health and Human Development, Bethesda,<br>USA) |
| Genetic reagent<br>(*D. rerio*) | *Tg(fli1aep:ERK-KTR-*<br>*Clover)<sup>uq39bh</sup>* | This study |  | Ben M Hogan (Organogenesis and Cancer<br>Program, Peter MacCallum Cancer Centre,<br>Australia) |
| Genetic reagent<br>(*D. rerio*) | *Tg(ubb:Mmu.Elk1-KTR-*<br>*mClover)<sup>vi1</sup>* | (Mayr et al.,<br>2018) | ZFIN ID: ZDB-ALT-190211-6 | Martin Distel<br>(Children's Cancer Research Institute, Austria) |
| Genetic reagent<br>(*D. rerio*) | *Tg(actb2:GCaMP6f)<sup>zf3076</sup>* | (Herzog et al.,<br>2019) | ZFIN ID: ZDB-ALT-200610-2 | Leah Herrgen<br>(Centre for Discovery Brain Sciences, University<br>of Edinburgh, Germany) |
| Genetic reagent<br>(*D.rerio*) | *Tg(kdrl:mCherry-CAAX)<sup>y171</sup>* | (Fujita et al., 2011) | ZFIN ID: ZDB-ALT-110429-3 | Brent M Weinstein (National Institute of Child Health and Human Development, Bethesda, USA) |
| Genetic reagent<br>(*D.rerio*) | *Tg(mpeg1:mCherry)<sup>gl23</sup>* | (Ellett et al., 2011) | ZFIN ID: ZDB-ALT-120117-2 | Graham Lieschke (Australian Regenerative Medicine Institute, Monash University, Australia) |
| Genetic reagent<br>(*D.rerio*) | *Tg(kdrl:EGFP)<sup>s843</sup>* | (Beis et al., 2005) | ZFIN ID: ZDB-ALT-050916-14 | Didier Stainier<br>(Max Planck Institute for Heart and Lung Research, Germany) |
| Sequence-based reagent | MO1-spi1b | (Rhodes et al., 2005) | ZFIN ID: ZDB-MRPHLNO-050224-1 | Genetools, LLC, OR, USA |
| Sequence-based reagent | MO3-csf3r | (Ellett et al., 2011) | ZFIN ID: ZDB-MRPHLNO-111213-1 | Genetools, LLC, OR, USA |
| Software, algorithm | FIJI | ImageJ<br>(http://imagej.nih.gov/ij/) |  | Image processing and analysis, Version Fiji version 1 |
| Software, algorithm | Imaris x64 | Bitplane, Belfast, UK |  | Image processing and analysis, Version 9.5.1 |
| Software, algorithm | GraphPad Prism | GraphPad Prism<br>(http://graphpad.com) |  | Statistics, Prism8: Version 8.3.0 |
| Software,<br>algorithm | R/R Studio | R project (r-<br>project.org) |  | Statistics, R version 4.0.2 |
| Chemical<br>compound, drug | SL327 (MEK signalling<br>inhibitor) | Merck,<br>Darmstadt,<br>Germany | S4069 | Diluted in DMSO |
| Chemical<br>compound, drug | Trametinib (MEK<br>signalling inhibitor) | Selleck<br>chemicals, TX,<br>USA | S2673 | Diluted in DMSO |
| Chemical<br>compound, drug | SU5416 | Merck,<br>Darmstadt,<br>Germany | S8442 | Diluted in DMSO |
| Chemical<br>compound, drug | AV951 | Adooq<br>Bioscience, CA,<br>USA | 475108-18-0 | Diluted in DMSO |
| Chemical<br>compound, drug | Nifedipine | Bio-Techne, MN,<br>USA | 1075 | Diluted in DMSO |
| Chemical<br>compound, drug | Amlodipine | Merck,<br>Darmstadt,<br>Germany | A5605 | Diluted in DMSO |

### Zebrafish

All zebrafish work was conducted in accordance with the guidelines of the animal ethics committees at the University of Queensland, Peter MacCallum Cancer Centre, University of Bristol, and the Children’s Cancer Research Institute. The transgenic zebrafish lines used were published previously as following: *Tg(fli1a:H2B-mCherry)^uq37bh^* (Baek et al., 2019), *Tg(fli1a:EGFP)^y1^* (Lawson and Weinstein, 2002), *Tg(ubb:Mmu.Elk1-KTR-mClover)^vi1^* (Mayr et al., 2018), *Tg(actb2:GCaMP6f)^zf3076^* (Herzog et al., 2019), *Tg(kdrl:mCherry-CAAX)^y171^* (Fujita et al., 2011), *Tg(mpeg1:mCherry)^gl23^* (Ellett et al., 2011), and *Tg(kdrl:EGFP)^s843^* (Beis et al., 2005). The *Tg(fli1aep:ERK-KTR-Clover)^uq39bh^* transgenic line (referred to as *Tg(fli1aep:EKC*) in this study) was generated for this study using Gateway cloning and transgenesis. The pENTR-ERKKTRClover plasmid (#59138) was purchased from Addgene.

### Live imaging and laser-inflicted vessel/tissue wounding

Embryos/Larvae at indicated stages were immobilised with Tricaine (0.08 mg/ml) and mounted laterally in either 1.2% ultra-low gelling agarose (specifically for **Video 6**), 0.25% low melting agarose (specifically for **Videos 7 and 8**, and **Figure 6A**), or 0.5% low melting agarose (Merck, Darmstadt, Germany, A9414-100G) as previously described (Okuda et al., 2018). Images were taken at indicated timepoints/timeframe using either a Zeiss LSM 710 confocal microscope (specifically for **Figure 1B-E**), Leica SP8 X WLL confocal microscope (specifically for **Video 6**), Leica TCS SP8 multiphoton microscope (specifically for **Videos 7 and 8**, and **Figure 6A**), Olympus Yokogawa CSU-W1 Spinning Disc Confocal microscope (specifically for **Figure 6-figure supplement 1K-T’**), or an Andor Dragonfly Spinning Disc Confocal microscope.

Muscle wounding in 30 hpf *Tg(ubb:Mmu.Elk1-KTR-mClover)* embryos were conducted as previously described (specifically for **Video 6**) (Mayr et al., 2018). Briefly, a laser-inflicted wound was introduced on mounted embryos using the Leica SP8 X FRAP module with the UV laser line of 405 nm at 85% laser power. Vessel wounding in 4 dpf *Tg(actb2:GCaMP6f);Tg(kdrl:mCherry-CAAX)* larvae were conducted as previously described (specifically for **Video 7 and 8**, and **Figure 6A**) (Gurevich et al., 2018). Briefly, a laser-inflicted wound was introduced on mounted larvae using a Micropoint laser (Spectra-Physics, CA, USA) connected to a Zeiss Axioplan II microscope with a laser pulse at a wavelength of 435 nm. All other tissue/vessel wounding in either 3 dpf (specifically for **Figure 3-figure supplement 1B,C,L-S**) or 4 dpf EC-EKC/mCherry or *Tg(kdrl:EGFP);Tg(mpeg1:mCherry)* larvae were conducted using either a Zeiss LSM 710 confocal microscope or a Olympus FVMPE-RS multiphoton microscope. Briefly, a laser-inflicted wound was introduced on mounted larvae using a two-photon laser at 790 nm (Zeiss LSM 710 confocal microscope) or 800 nm (Olympus FVMPE-RS multiphoton microscope) at 80% laser power (Mai Tai, Spectra-Physics, CA, USA). The area of laser ablation for vessel wounding experiments was made consistent for all experiments (height: 40 μm, width: 15 μm). All vessel wounding was conducted on the ISV dorsal to the cloaca.

For **Video 1**, time-lapse images of ISVs in 24-25 EC-EKC/mCherry embryos were acquired every 14-17 seconds for 40 minutes using an Andor Dragonfly Spinning Disc Confocal microscope. Difference in time intervals were due to difference in z section number in different embryos. Pre-division ISV tip ECs with cytoplasmic H2B-mCherry localisation were selected for imaging. For **Videos 3-5**, time-lapse images of ISVs in 4 dpf EC-EKC/mCherry larvae were taken every minute for 20 minutes using an Andor Dragonfly Spinning Disc Confocal microscope, wounded as described above using a Zeiss LSM 710 confocal microscope, transferred to an Andor Dragonfly Spinning Disc Confocal microscope (allowing for 2 minutes to transfer the larvae and initiate imaging) and re-imaged every minute for another 20 minutes. As a control (**Video 2**), time-lapse images of ISVs in 4 dpf EC-EKC/mCherry larvae were taken every minute for 41 minutes. For **Video 6**, time-lapse images of the trunk in a 30 hpf *Tg(ubb:Mmu.Elk1-KTR-mCherry)* embryo were acquired every 21 minutes from 5 mpa until 3 hpa using a Leica SP8 X WLL confocal microscope. For **Video 8**, time-lapse images of ISVs in 4 dpf *Tg(actb2:GCaMP6f);Tg(kdrl:mCherry-CAAX)* larvae were acquired every minute from 5 mpa until 20 mpa using a Leica SP8 confocal microscope. As a control (**Video 7**), time-lapse images of ISVs in 4 dpf *Tg(actb2:GCaMP6f);Tg(kdrl:mCherry-CAAX)* larvae were acquired every minute for 15 minutes using a Leica SP8 confocal microscope.

### Morpholino injections

The *spi1b* and *csf3r* morpholinos used in this study have been validated and described previously (Rhodes et al., 2005;Ellett et al., 2011;Pase et al., 2012). A cocktail of *spi1b* (5ng) and *csf3r* (2.5ng) morpholinos were injected into 1-4 cell stage EC-EKC/mCherry or *Tg(mpeg1:mCherry)* embryos as previously described (Pase et al., 2012). ISVs of 3 dpf morphants/uninjected controls were imaged before vessel wounding, wounded as described above, and reimaged at 15 mpa. Non-ablated 3 dpf EC-EKC/mCherry morphants/uninjected controls were imaged, and re-imaged 15 minutes later. Macrophage numbers (*mpeg1:mCherry*-positive) in 3 dpf embryos (**Figure 3-figure supplement 1E,F**) or 4 dpf larvae (**Figure 3-figure supplement 1A-C**) were manually quantified using the cell counter tool in FIJI.

### Drug treatments

For investigating Erk activity in ISV tip ECs in 28 hpf embryos following drug treatment, 27 hpf *Tg(fli1aep:EKC);Tg(fli1a:H2B-mCherry)* embryos were treated for an hour with either 0.5% DMSO (vehicle control), 15 μM SL327, 4 μM SU5416, or 500 nM AV951 diluted in E3 medium with 0.003% 1-phenyl-2-thiourea (PTU) and imaged as described above at 28 hpf. Up to 5 ISV tip ECs were quantified per embryo. For investigating the role of prolonged EC Erk activity in vessel regeneration, ISVs of 4 dpf *Tg(fli1aep:EKC)* larvae were wounded as described above and were treated with either 0.5% DMSO (vehicle control), 4 μM SU5416, 15 μM SL327, or 1 μM Trametinib for 24 hours and imaged as described above at 5 dpf (24 hpa). For measuring Erk activity in ECs pre- and post-ablation in 4 dpf larvae following drug treatment, 4 dpf EC-EKC/mCherry larvae were first treated for an hour with either 0.5% DMSO, 15 μM SL327, 4 μM or 10 μM SU5416, or 500 nM AV951. ISVs of these larvae were imaged then wounded as described above in the presence of respective drugs at indicated concentrations in the mounting media. The same larvae were reimaged at 15 mpa. Alternatively, larvae were removed from mounting media following vessel wounding and incubated in respective drugs at indicated concentrations in E3 media, before being remounted and imaged at 3 hpa.

For Nefidipine and Amlopidine treatment, 4 dpf EC-EKC/mCherry larvae were first treated for 30 minutes with either 1% DMSO, 50 μM Nifedipine, or 100 μM Amlodipine. This was because treatment for 1 hour with either 50 μM nifedipine or 100 μM Amlodipine resulted in mortalities due to reduced cardiac function. The ISVs of these larvae were imaged and wounded as described above and reimaged 15 mpa. Alternatively, 4 dpf EC-EKC/mCherry larvae were imaged before vessel wounding, and removed from mounting media following vessel wounding and incubated in 1% DMSO. 30 minutes before 3 hpa, larvae were treated with 50 μM Nifedipine or continued its treatment with 1% DMSO, before being remounted in the presence of respective drugs at indicated concentrations and reimaged 3 hpa. Non-ablated 4 dpf EC-EKC/mCherry larvae controls were imaged, then reimaged either 15 mpa or 3 hpa as described above.

### Image processing and analysis

Images were processed with image processing software FIJI version 1 (Schindelin et al., 2012) and Imaris x64 (Bitplane, Version 9.5.1). Erk activity in ECs were measured by either comparing nuclear/cytoplasm EKC intensity, nuclear EKC/H2B- mCherry intensity, or nuclear EKC intensity. The nuclear/cytoplasm EKC intensity was quantified as described before (Kudo et al., 2018) with modifications, using a semi-autonomous custom written script in the ImageJ macro language. Briefly, z stack images were first processed into a maximum intensity z-projection. H2B- mCherry-positive EC nuclei underwent thresholding and were selected as individual regions of interest (ROI). The EKC channel was converted to a 32-bit image with background (non-cell associated) pixels converted to NaN. The average pixel intensity of EKC in the nuclei ROIs were measured (nuclear EKC intensity). Nuclei ROIs were then expanded and converted to a banded selection of the adjacent cytoplasmic area and the average pixel intensity of EKC within the expanded ROIs were measured (cytoplasm EKC intensity). The custom written ImageJ macro is available here: [https://github.com/NickCondon/Nuclei-Cyto_MeasuringScipt].

The average pixel intensity of either nuclear EKC or H2B-mCherry of ECs in 3D was quantified using Imaris software. The entire EC nucleus was masked using the H2B- mCherry signal. **Figures 1J and K** represent averages of data within each minute. For Embryos/larvae exposed to long-term time-lapse (for example **Videos 2-5**), or ablated with high-powered multiphoton laser for ablation studies, difference in photostability between fluorophores could significantly alter the ratio of nuclear EKC/H2B-mCherry intensity (Lam et al., 2012). Therefore, we either compared the ratio of nuclear EKC intensities between ECs within the same fish (for example **Video 1**), or we normalised EC nuclear EKC intensity with the average EKC intensity of another EKC-expressing structure (for example **Videos 2-5**). For larvae that underwent laser-inflicted wounding, nuclear EKC intensity pre- and post-ablation was normalised with the average pixel intensity of EKC of the entire DA within 2 somite length. The ROI that covers the same DA region in pre- and post-wounded larvae was manually selected on a maximum intensity z-projection of the EKC channel, and average pixel intensity was calculated using FIJI. Datasets were presented as either the ratio of post/pre-ablation normalised nuclear EKC intensity, or as normalised nuclear EKC intensity further normalised to normalised nuclear EKC intensity in 2 mpa ECs (specifically for **Figure 6O**). 3 closest ECs from the wounded site in both ablated and adjacent ISVs were quantified, except for **Figures 5K and 6G**, where 5 closest ECs from the wounded site in ablated ISVs were analysed. For **Videos 2-5,** reduction in EKC intensity due to photobleaching was minimised using the bleach correction tool (correction method: Histogram Matching) in FIJI, however quantifications were all done using raw data.

GCaMP6f average pixel intensity on ISVs and unablated tissue in 4 dpf *Tg(actb2:GCaMP6f);Tg(kdrl:mCherry-CAAX)* larvae was measured using FIJI. Maximum intensity z-projection images of both GCaMP6f and mCherry-CAAX channels were first corrected for any drift in x/y dimensions. A ROI was drawn around the mCherry-CAAX-positive ISV segment nearest to the site of injury (an area consistently between 100-150 μm^2^) and the average pixel intensity of GCaMP6f within the ROI at each timepoints were measured using FIJI. Similar measurements were acquired for adjacent ISVs, ISVs in unablated control larvae, and uninjured tissue, maintaining consistent size of ROI within each biological replicate. ISV GCaMP6f average pixel intensity was normalised to the average pixel intensity in uninjured tissue GCaMP6f within the same larvae.

The percentage of ISV height was measured by dividing the total horizontal height of the ISV with the prospective total horizontal height of the ISV (the horizontal height from the base ISV/DA intersection to the tip of the ISV/DLAV intersection. Ellipticity (oblate) of ISV tip ECs were quantified using Imaris software.

### Statistics

Graphic representations of data and statistical analysis was performed using either Prism 8 Version 8.3.0 or R software. Mann-Whitney test was conducted when comparing two datasets and Kruskal-Wallis test was conducted when comparing multiple datasets using Prism 8. Natural permutation test was used to test for differences between the population mean curve for datasets in **Figure 3E,F** and **Figure 4C,D** using R software. Stars indicate p-value as level of significance with p≤0.001 (***), p≤0.01 (**), p≤0.05 (*), and p>0.05 (not significant, n.s.). Error bars in all graphs represent standard deviation.

## Acknowledgements

This work was supported by NHMRC grants 1164734 and 1165117. BMH was supported by an NHMRC fellowship 1155221. We thank Dr Enid Lam and Dr Elizabeth Mason for their technical assistance. Imaging was performed in the Australian Cancer Research Foundation’s Cancer Ultrastructure and Function Facility at IMB, Centre for Advanced Histology and Microscopy at Peter MacCallum Cancer Centre, Wolfson Bioimaging Facility at University of Bristol, and the Zebrafish platform Austria for preclinical drug screening at the Children’s Cancer Research Institute supported by the Austrian Research Promotion Agency (FFG) project 7640628 (Danio4Can).

## Author contributions

Kazuhide S. Okuda, Conceptualization, Formal analysis, Investigation, Methodology, Writing-original draft; Mikaela Keyser, Formal analysis, Investigation; David B. Gurevich, Formal analysis, Investigation; Caterina Sturtzel, Investigation; Scott Patterson, Investigation; Huijun Chen, Investigation; Mark Scott, Methodology; Nicholas Condon, Methodology; Paul Martin, Resources, Supervision; Martin Distel, Conceptualization, Resources, Supervision; Benjamin M. Hogan, Conceptualization, Resources, Supervision, Funding acquisition, Writing-review and editing.

## Supplementary figure legends

**Figure 1-figure supplement 1:**
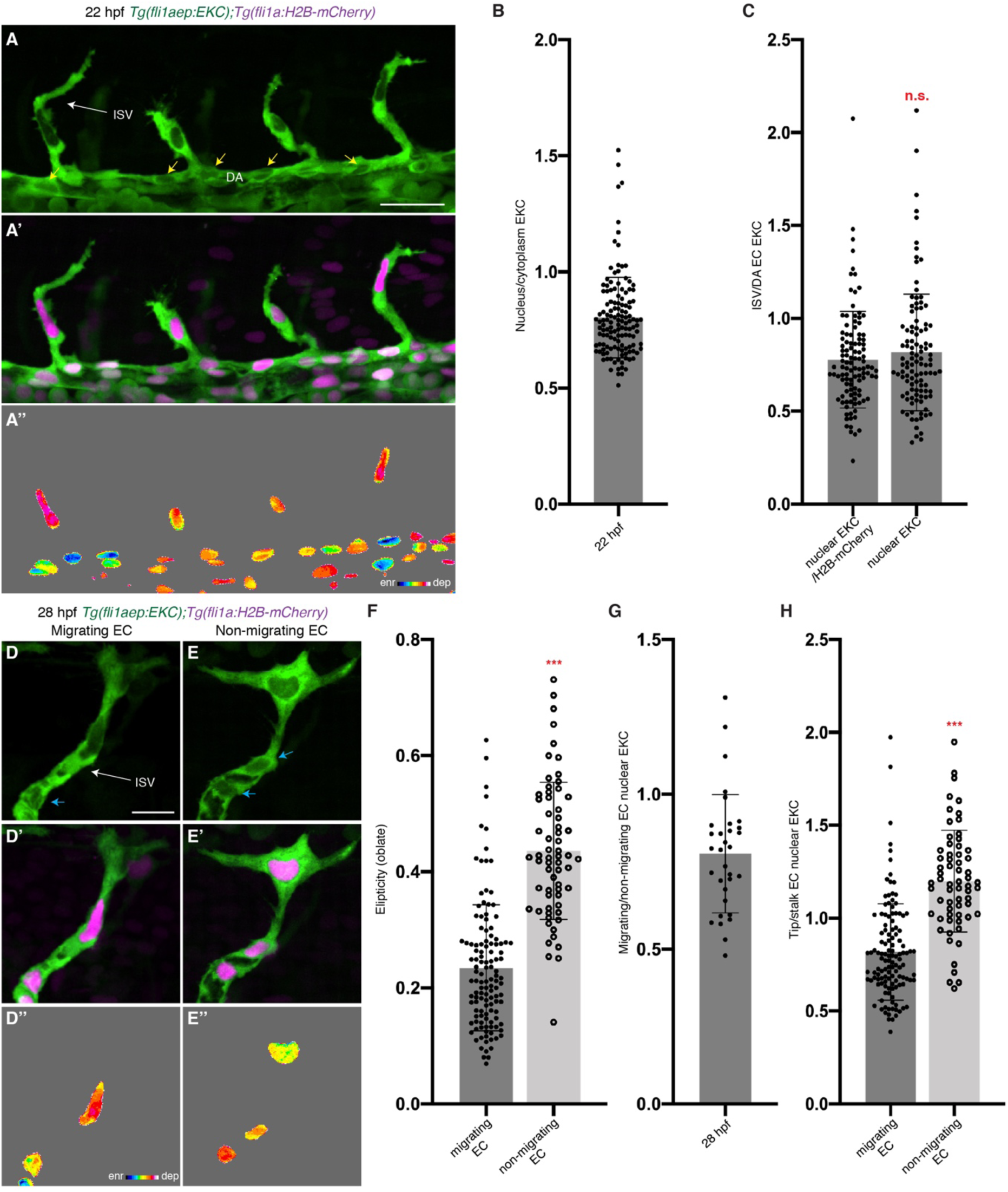
The *Tg(fli1aep:EKC)* transgenic line reports physiologically relevant Erk-signalling during primary angiogenesis. (**A-A’’**) High Erk-activity in tip-cells. Lateral spinning disc confocal images of budding ISVs in a 22 hpf *Tg(fli1ep:EKC);Tg(fli1a:H2B-mCherry)* embryo. Image A shows the *fli1aep:EKC* expression, image A’ shows both the *fli1ep:EKC* and the *fli1a:H2B-mCherry* expression, while image A’’ shows the nuclear *fli1aep:EKC* expression with intensity difference represented in 16 colour LUT (Fiji). The *fli1a:H2B-mCherry* signal was used to mask the nucleus. Yellow arrows point to DA ECs with nuclear depleted EKC localisation. (B) Quantification of nucleus/cytoplasm EKC intensity in sprouting ISV ECs of 22 hpf embryos (0.803, 133 ECs, n=37 embryos). (C) Quantification of sprouting ISV EC/ DA “stalk” EC nuclear EKC intensity in 22 hpf embryos (109 ECs, n=37 embryos). DA ECs closest to the sprouting ISV ECs were quantified. Both ratios of nuclear EKC/H2B-mCherry intensity (0.777) and nuclear EKC alone (0.817) show higher Erk activity in sprouting ISV ECs when compared to DA “stalk” ECs. (**D-E’’**) Nuclear elongation and Erk-activity correlate. Lateral spinning disc confocal images of either an ISV with migrating tip EC (D) or an ISV with non-migrating tip EC (E) in 28 hpf *Tg(fli1aep:EKC);Tg(fli1a:H2B-mCherry)* embryos. Images D and E show the *fli1aep:EKC* expression, images D’ and E’ show both *fli1ep:EKC* and *fli1a:H2B-mCherry* expressions, while images D’’ and E’’ show nuclear *fli1aep:EKC* expression with intensity difference represented in 16 colour LUT (Fiji). The *fli1a:H2B-mCherry* signal was used to mask the nucleus. Light blue arrow show ISV stalk ECs with nuclear depleted EKC localisation. (F) Quantification of EC ellipticity (oblate) in migrating (124 ECs, n=45 embryos) and non-migrating ISV tip ECs (64 ECs, n=35 embryos) in 28 hpf embryos. (G) Quantification of migrating/non-migrating ISV tip EC nuclear EKC intensity in 28 hpf embryos (0.808, n=32 embryos). Only embryos with both migrating/non-migrating ISV tip ECs in the same image were quantified. (H) Quantification of tip/stalk ISV EC nuclear EKC intensity of either migrating tip ECs or non-migrating tip ECs in 28 hpf embryos (124 migrating tip ECs, n=45 embryos, 64 non-migrating tip ECs, n=35 embryos). ISV: intersegmental vessel; DA: dorsal aorta. Statistical test: Mann-Whitney test was conducted for graphs C, F, and H. Error bars represent standard deviation. Scale bars: 25 μm for image A, 15 μm for image D.

**Figure 3-figure supplement 1:**
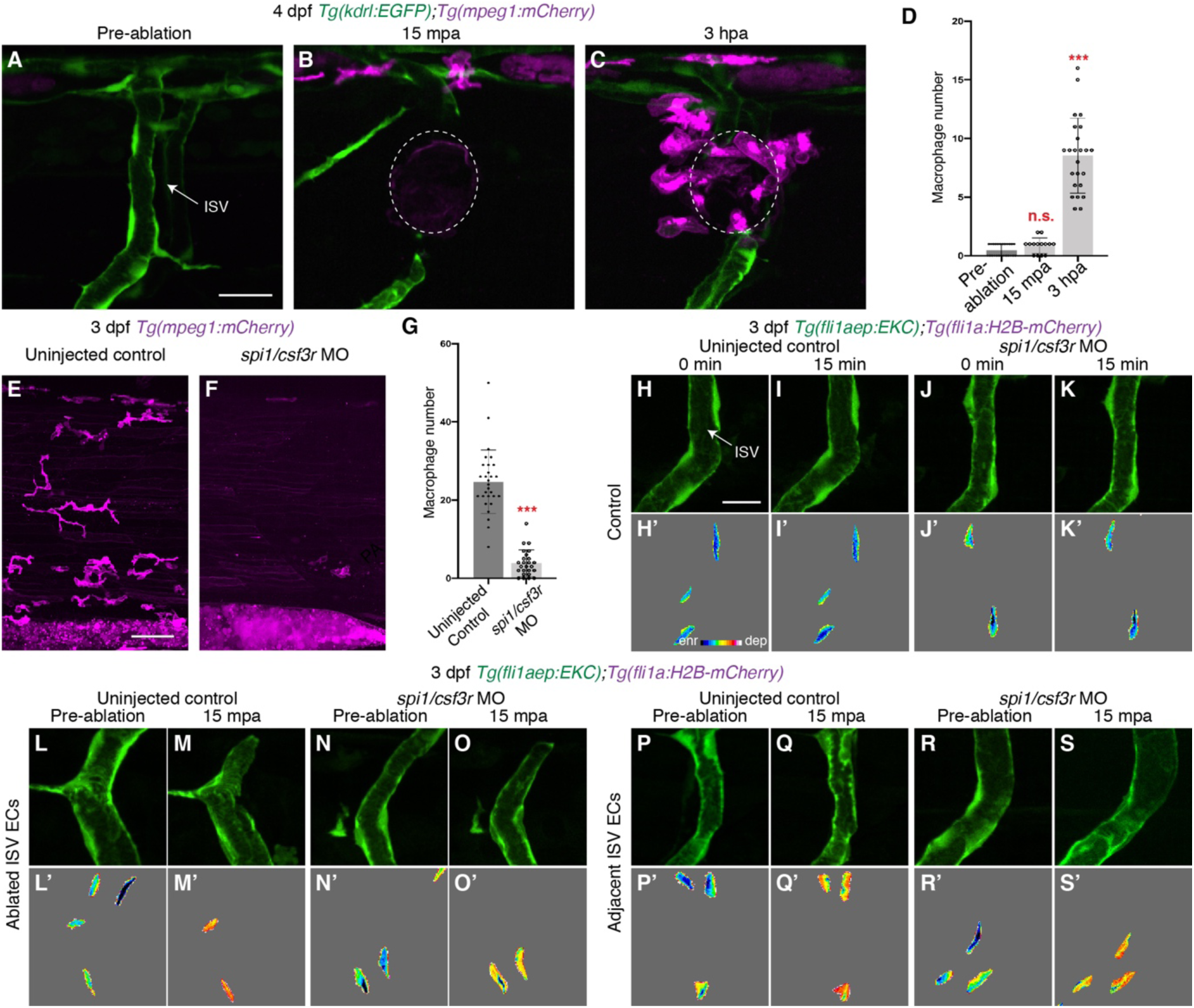
Macrophages are not required for rapid Erk activation following vessel wounding. (**A-C**) Lateral spinning disc confocal images of 4 dpf *Tg(kdrl:EGFP);Tg(mpeg1:mCherry)* larvae at either pre-ablation (A), 15 mpa (B), or 3 hpa (C). White circles show the wounded site of each larvae. (**D**) Quantification of macrophage number recruited to the wounded site pre-ablation (n=25 embryos), 15 mpa (n=14 embryos) or 3 hpa (n=24 embryos). (**E,F**) Lateral spinning disc confocal images of 3 dpf *Tg(mpeg1:mCherry)* uninjected control (E) or *spi1*/*csf3r* morphants (F). (**G**) Quantification of macrophage number within the trunk in 3 somites length in 3 dpf *Tg(mpeg1:mCherry)* uninjected control (n=29 embryos) or *spi1*/*csf3r* morphants (n=25 embryos). (**H-S’**) Lateral spinning disc confocal images of ISV ECs in 3 dpf *Tg(fli1aep:EKC);Tg(fli1a:H2B-mCherry)* uninjected control (H-I’, L-M’, P-Q’) and *spi/csf3r* morphants (J-K’, N-O’, R-S’). Images H-K’ show non ablated control ISV ECs, images L-O’ show ablated ISV ECs, and images P-S’ show adjacent ISV ECs. Images L,N,P,R were taken pre-ablation, while images M,O,Q,S were taken 15 mpa. Images H-S show the *fli1aep:EKC* expression, and images H’-S’ shows the nuclear *fli1aep:EKC* expression with intensity difference represented in 16 colour LUT (Fiji). The *fli1a:H2B-mCherry* signal was used to mask the nucleus. ISV: intersegmental vessel; Statistical test: Kruskal Wallis test was conducted for graph D and Mann-Whitney test was conducted for graph G. n.s. represents not significant. Error bars represent standard deviation. Scale bars: 20 μm for image A, 50 μm for image E, 15 μm for image H.

**Figure 3-figure supplement 2:**
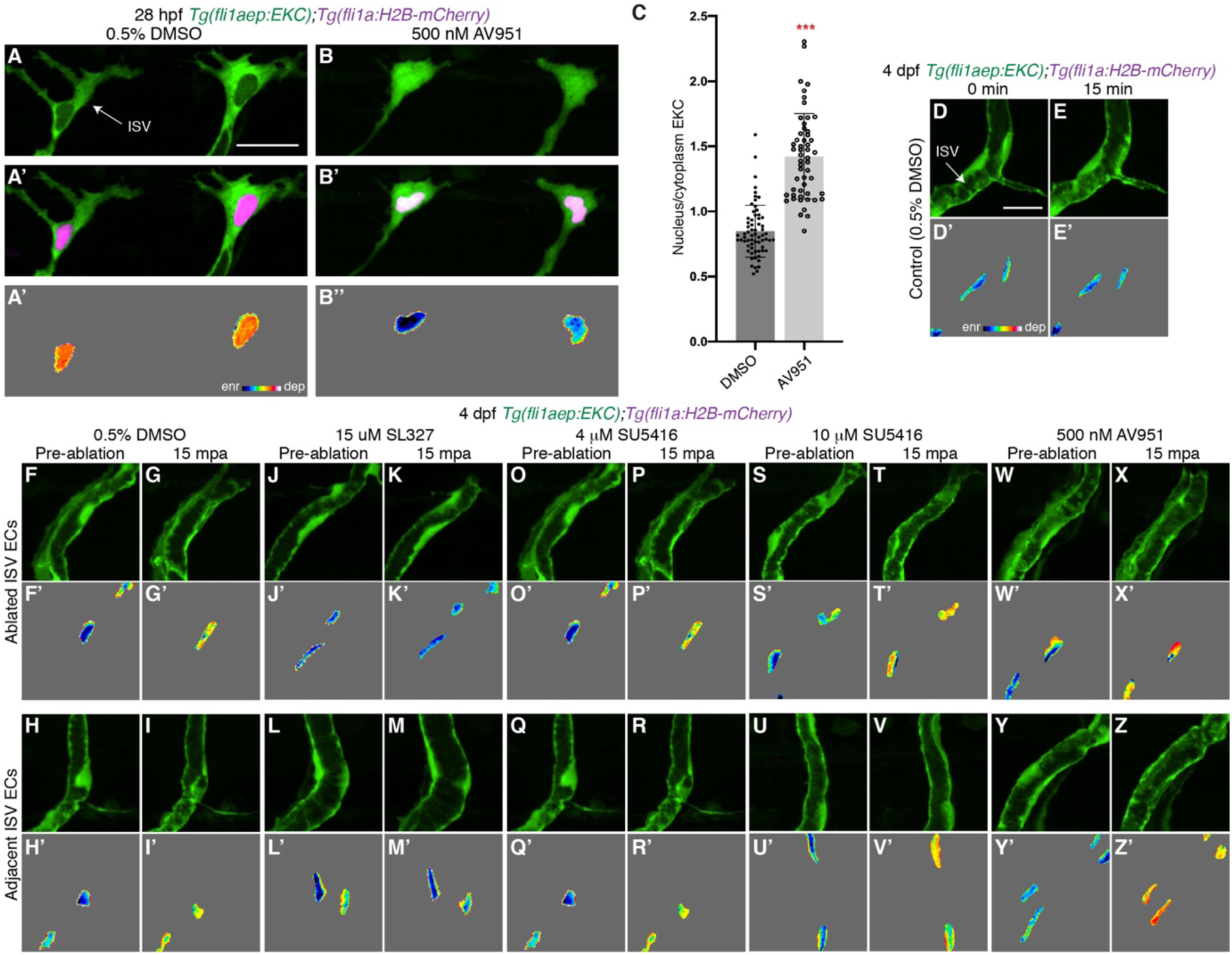
Vegfr-signalling is not required for rapid Erk activation following vessel wounding. (**A**-**B’’**) Lateral spinning disc confocal images of ISV ECs in 28 hpf *Tg(fli1aep:EKC);Tg(fli1a:H2B-mCherry)* embryos treated for an hour with either 0.5% DMSO (A-A’’) or 500 nM AV951 (B-B’’). Images A and B show the *fli1aep:EKC* expression, while images A’ and B’ show both the *fli1aep:EKC* and the *fli1a:H2B-mCherry* expression. Images A’’ and B’’ show the nuclear *fli1aep:EKC* expression with intensity difference represented in 16 colour LUT (Fiji). The *fli1a:H2B-mCherry* signal was used to mask the nucleus. (**C**) Quantification of nucleus/cytoplasm EKC intensity in ISV tip ECs of 28 hpf embryos treated with either 0.5% DMSO (0.849, 65 ECs, n=14 embryos) or 500 nM AV951 (1.423, 53 ECs, n=12 embryos). (**D-Z’**) Lateral spinning disc confocal images of ISV ECs in 4 dpf *Tg(fli1aep:EKC);Tg(fli1a:H2B-mCherry)* larvae treated with either 0.5% DMSO (D-I’), 15 μM SL327 (J-M’), 4 μM SU5416 (O-R’), 10 μM SU5416 (S-V’), or 500 nM AV951 (W-Z’). Images D-E’ show non-ablated control ISV ECs. Images F-G’, J-K’, O-P’, S-T’ and W-X’ show ablated ISV ECs. Images H-I’, L-M’, Q-R’, U-V’ and Y-Z’ show adjacent ISV ECs. Images F,H,J,L,O,Q,S,U,W,Y were taken pre-ablation and images G,I,K,M,P,R,T,V,X,Z were taken 15 mpa. Images D-Z show the *fli1aep:EKC* expression, and images D’-Z’ show the nuclear *fli1aep:EKC* expression with intensity difference represented in 16 colour LUT (Fiji). The *fli1a:H2B-mCherry* signal was used to mask the nucleus. ISV: intersegmental vessel; Statistical test: Mann-Whitney test was conducted for graph C. Error bars represent standard deviation. Scale bars: 25 μm for image A, 15 μm for image D.

**Figure 4-figure supplement 1:**
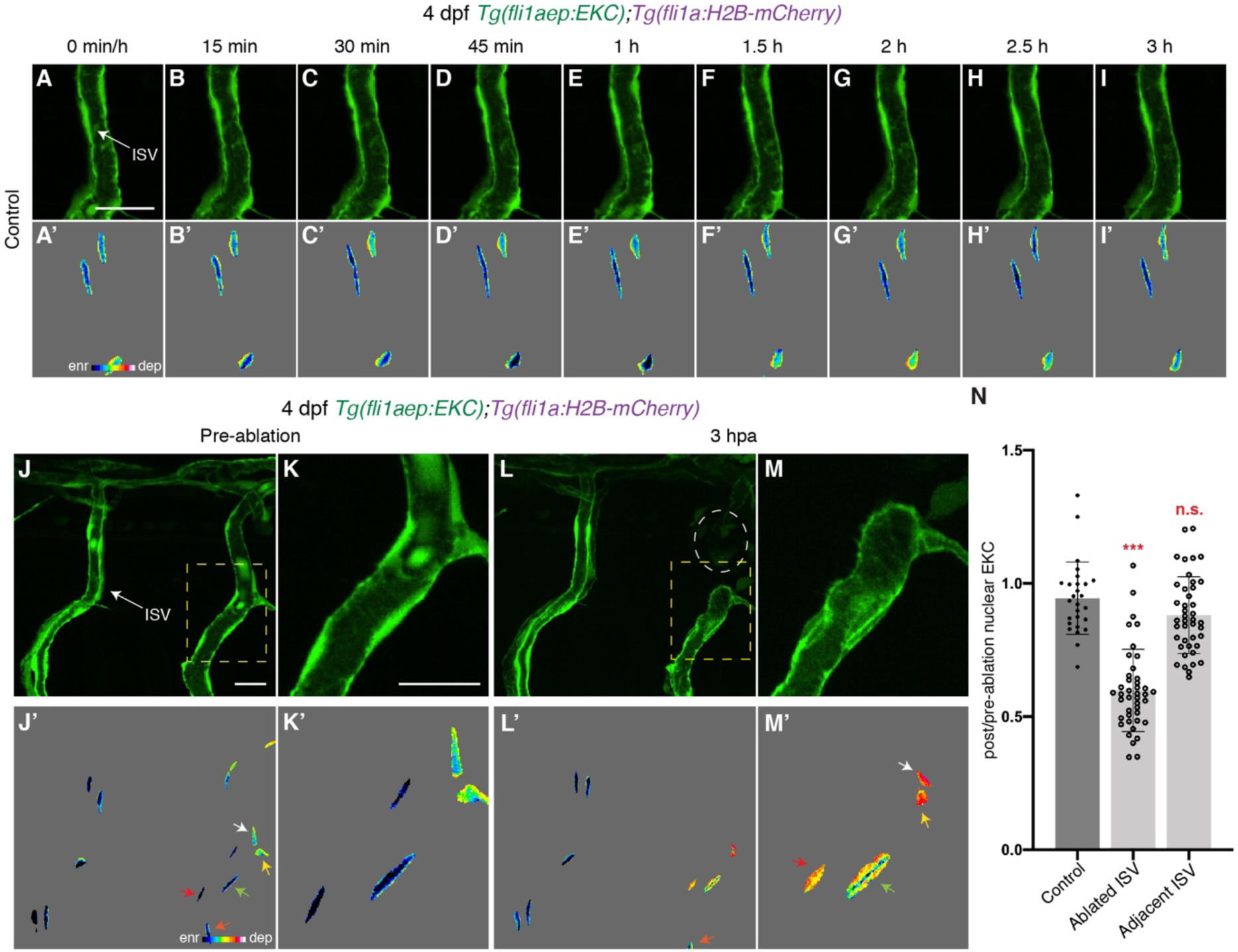
Distinct Erk activity between ablated and adjacent ISV ECs 3 hpa. (**A-I’**) Lateral spinning disc confocal images of ISV ECs in 4 dpf *Tg(fli1aep:EKC);Tg(fli1a:H2B-mCherry)* larvae at indicated timepoints. Images A-I show the *fli1aep:EKC* expression, while images A’-I’ show the nuclear *fli1aep:EKC* expression with intensity difference represented in 16 colour LUT (Fiji). The *fli1a:H2B-mCherry* signal was used to mask the nucleus. (**J-M’**) Lateral spinning disc confocal images of ablated and adjacent ISV ECs in 4 dpf *Tg(fli1aep:EKC);Tg(fli1a:H2B-mCherry)* larvae before (J-K’), and 3 hours following vessel wounding (L-M’). Images J-M show the *fli1aep:EKC* expression, while images J’-M’ show the nuclear *fli1aep:EKC* expression with intensity difference represented in 16 colour LUT (Fiji). The *fli1a:H2B-mCherry* signal was used to mask the nucleus. Images K and M are higher magnification images of the yellow boxes in images J and L. White circle in image L show the wounded site. Arrows indicate first (white), second (yellow), third (green), forth (red), and fifth (orange) ECs from the wounded site. (**N**) Quantification of post/pre-ablation nuclear EKC intensity of ECs in non-ablated control ISVs (27 ECs, n=9 larvae), ablated ISVs (42 ECs, n=14 larvae), and adjacent ISVs (42 ECs, n=14 larvae) 3 hpa. ISV: intersegmental vessel; Statistical test: Kruskal Wallis test was conducted for graph N. n.s. represents not significant. Error bars represent standard deviation. Scale bars: 20 μm for image A, 20 μm for images J and K.

**Figure 4-figure supplement 2:**
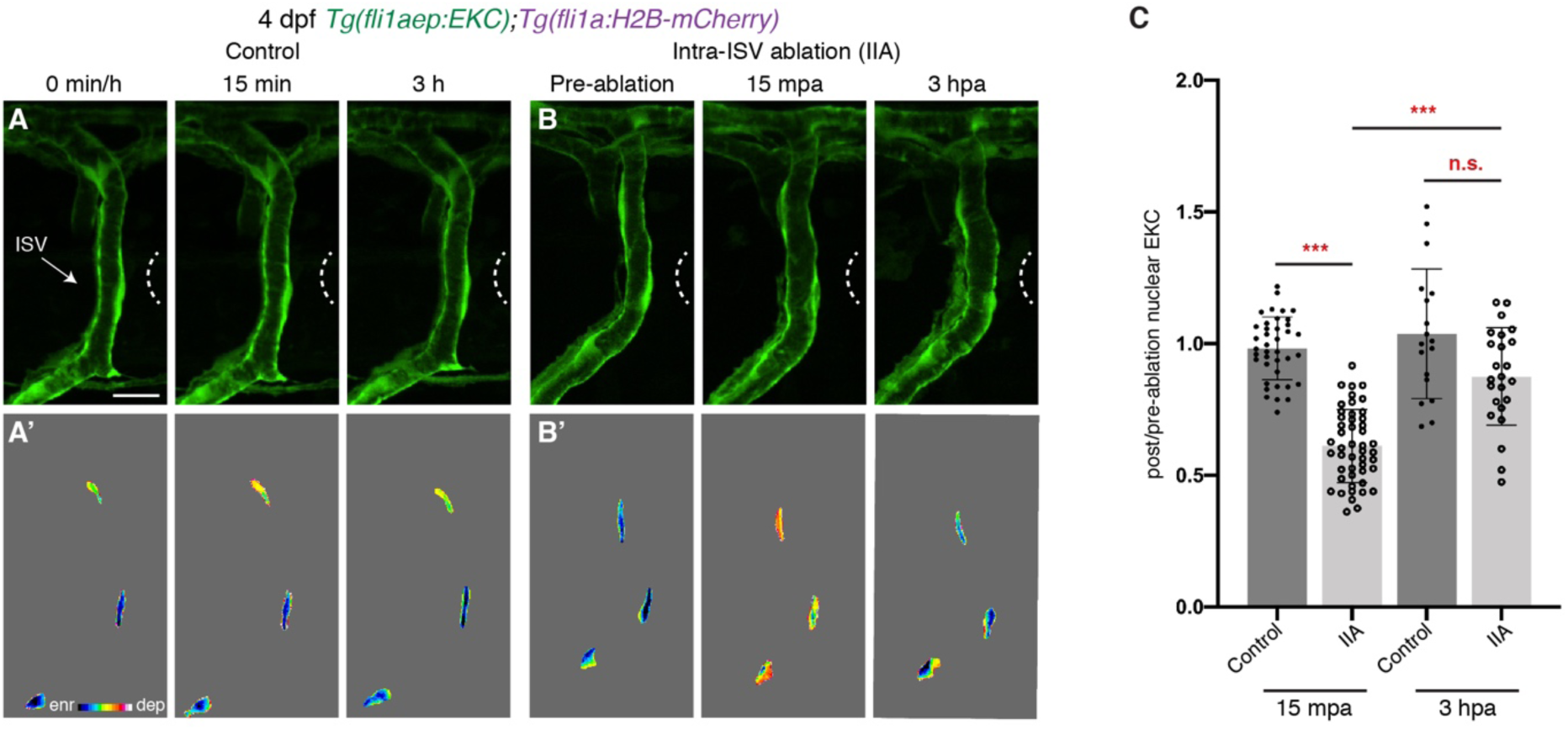
Vessel wounding is required for sustained Erk activity in ablated ISV ECs. (**A,B**) Lateral spinning disc confocal images of ISV ECs in 4 dpf *Tg(fli1aep:EKC);Tg(fli1a:H2B-mCherry)* larvae at 0min/pre-ablation (left), 15 minutes/15 mpa (middle), or 3 hours/3 hpa (right). Images A show ISVs in non-ablated control larvae, and images B show ISVs in larvae with tissue ablated in between two ISVs (Intra-ISV ablation (IIA)). Images A and B show the *fli1aep:EKC* expression, while images A’ and B’ show the nuclear *fli1aep:EKC* expression with intensity difference represented in 16 colour LUT (Fiji). The *fli1a:H2B-mCherry* signal was used to mask the nucleus. White dotted lines show the wounded sites. (**C**) Quantification of post/pre-ablation nuclear EKC intensity of ECs in either non-ablated control ISVs or Intra-ISV ablation ISVs at 15 mpa (control, 39 ECs, n=13 larvae; intra-ISV ablation, 48 ECs, n=16 larvae) or 3 hpa (control, 18 ECs, n=6 larvae; intra-ISV ablation, 24 ECs, n=8 larvae). ISV: intersegmental vessel; Statistical test: Kruskal Wallis test was conducted for graph C. n.s. represents not significant. Error bars represent standard deviation. Scale bar: 15 μm

**Figure 5-figure supplement 1:**
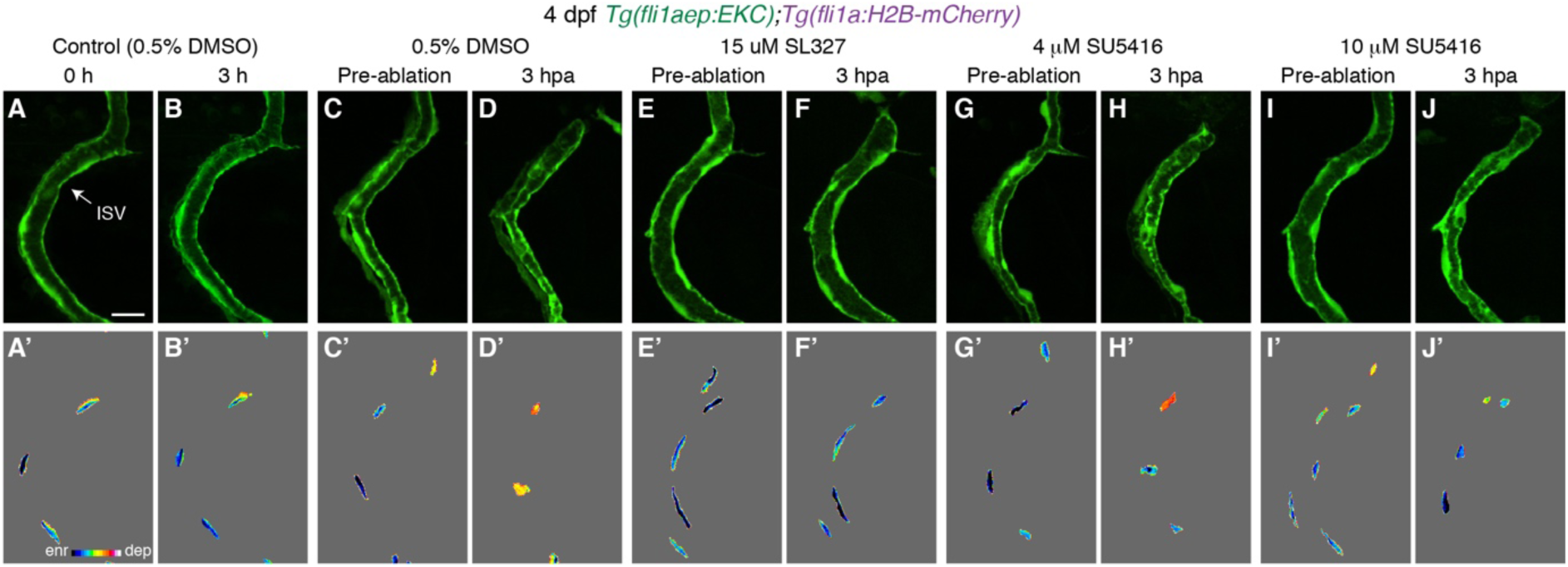
Vegfr-signalling is required to sustain Erk activity in ablated ISV ECs following vessel wounding. (**A-J’**) Lateral spinning disc confocal images of ISV ECs in 4 dpf *Tg(fli1aep:EKC);Tg(fli1a:H2B-mCherry)* larvae treated with either 0.5% DMSO (A-D’), 15 μM SL327 (E-F’), 4 μM SU5416 (G-H’), or 10 μM SU5416 (I-J’). Images A-B’ show non-ablated control ISV ECs. Images C,E,G,I were taken pre-ablation and images D,F,H,J were taken 3 hpa. Images A-J show the *fli1aep:EKC* expression, and images A’-J’ show the nuclear *fli1aep:EKC* expression with intensity difference represented in 16 colour LUT (Fiji). The *fli1a:H2B-mCherry* signal was used to mask the nucleus. ISV: intersegmental vessel; Scale bar: 15 μm

**Figure 6-figure supplement 1:**
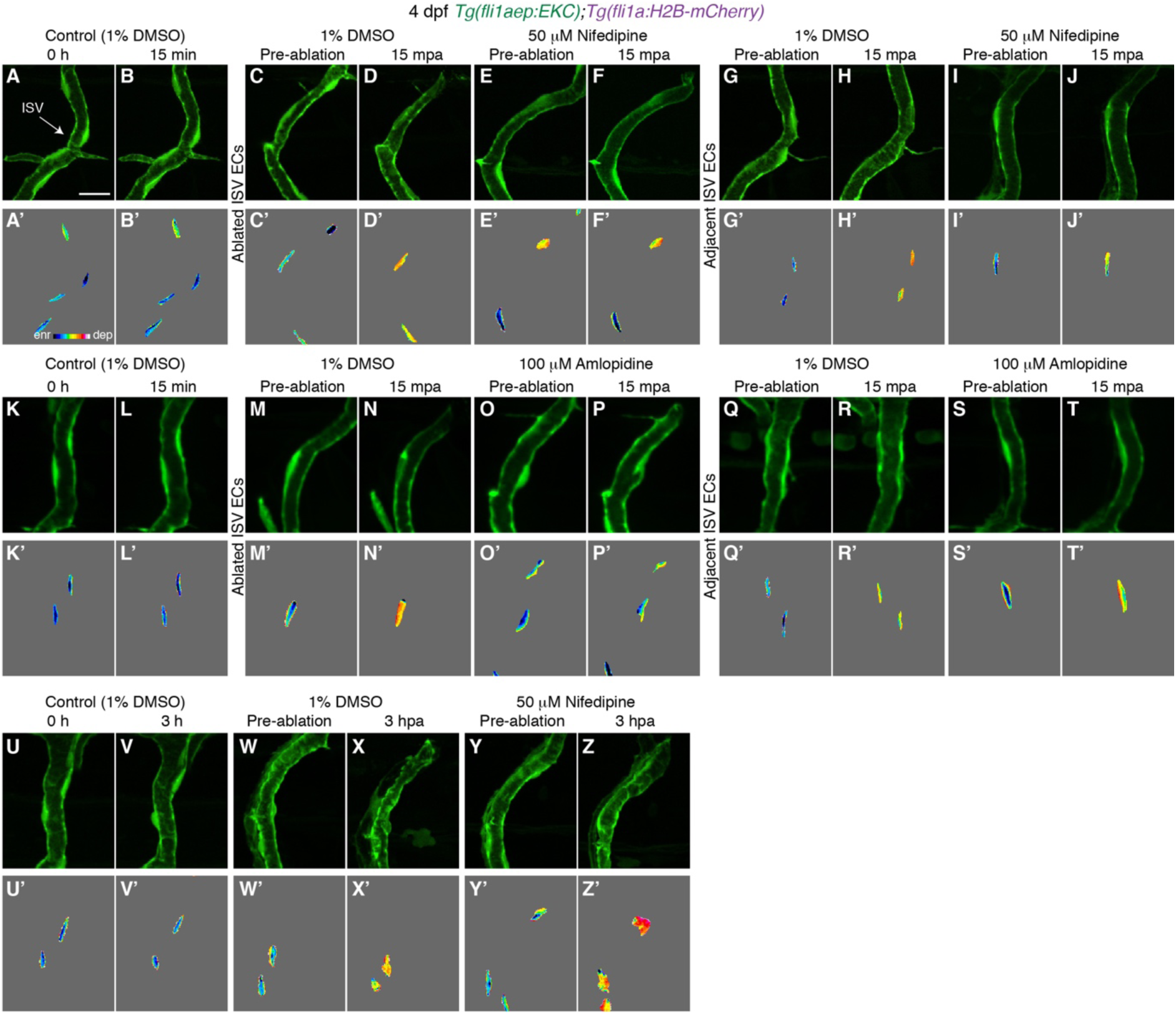
Ca^2+^ signalling is required for rapid Erk activation in ablated ISV ECs. (A-Z’) Lateral spinning disc confocal images of ISV ECs in 4 dpf *Tg(fli1aep:EKC);Tg(fli1a:H2B-mCherry)* larvae treated with either 1% DMSO (A-D’,G-H’,K-N’,Q-R’,U-X’), 50 μM Nifedipine (E-F’,I-J’,Y-Z’), or 100 μM Amlopidine (O-P’,S-T’). Images A-B’,K-L’,U-V’ show non-ablated control ISV ECs. Images C,E,G,I,M,O,Q,S,W,Y were taken pre-ablation, images D,F,H,J,N,P,R,T were taken 15 mpa, and images X and Z were taken 3 hpa. Images A-Z show the *fli1aep:EKC* expression, and images A-Z’ show the nuclear *fli1aep:EKC* expression with intensity difference represented in 16 colour LUT (Fiji). The *fli1a:H2B-mCherry* signal was used to mask the nucleus. ISV: intersegmental vessel. Scale bar: 15 μm

**<u>Video 1: ISV daughter ECs show asymmetric Erk activity following cytokinesis.</u>** Time-lapse video of a ISV tip EC undergoing mitosis in a 24-25 hpf *Tg(fli1aep:EKC);Tg(fli1a:H2B-mCherry)* embryo. Left panel shows the *fli1aep:EKC* expression, middle panel shows the *fli1a:H2B-mCherry* expression, and the right panel shows the nuclear *fli1aep:EKC* expression with intensity difference represented in 16 colour LUT (Fiji). The *fli1a:H2B-mCherry* signal was used to mask the nucleus. Z stacks were acquired every 15.5 seconds for 40 minutes using an Andor Dragonfly Spinning Disc Confocal microscope. Photobleaching was minimised using the bleach correction tool (correction method: Histogram Matching) in FIJI.

ISV: intersegmental vessel; DA: dorsal aorta. Scale bar: 25 μm.

**<u>Video 2: ISV ECs in 4 dpf larvae have minimal Erk activity.</u>** Time-lapse video of the trunk vessels in a 4 dpf *Tg(fli1aep:EKC);Tg(fli1a:H2B-mCherry)* larva at indicated timepoints. Left panel shows the *fli1aep:EKC* expression, middle panel shows both *fli1aep:EKC* and *fli1a:H2B-mCherry* expression, and the right panel shows the nuclear *fli1aep:EKC* expression with intensity difference represented in 16 colour LUT (Fiji). The *fli1a:H2B-mCherry* signal was used to mask the nucleus. Z stacks were acquired every minute for 41 minutes using an Andor Dragonfly Spinning Disc Confocal microscope. Photobleaching was minimised using the bleach correction tool (correction method: Histogram Matching) in FIJI.

ISV: intersegmental vessel; DA: dorsal aorta. Scale bar: 20 μm.

**<u>Video 3: Both ablated and adjacent ISV ECs rapidly activate Erk-signalling following vessel wounding.</u>** Time-lapse video of the trunk vessels in a 4 dpf *Tg(fli1aep:EKC);Tg(fli1a:H2B-mCherry)* larva before (pre-ablation) and after (post-ablation) vessel wounding at indicated timepoints. Post-ablation video starts at 2 minutes post-ablation due to the time taken to transfer the larvae between microscopes and for preparation of imaging. Left panel shows the *fli1aep:EKC* expression, middle panel shows both *fli1aep:EKC* and *fli1a:H2B-mCherry* expression, and the right panel shows the nuclear *fli1aep:EKC* expression with intensity difference represented in 16 colour LUT (Fiji). The *fli1a:H2B-mCherry* signal was used to mask the nucleus. Z stacks were acquired every 1 minutes for 20 minutes before and after vessel wounding using an Andor Dragonfly Spinning Disc Confocal microscope. Photobleaching was minimised using the bleach correction tool (correction method: Histogram Matching) in FIJI.

ISV: intersegmental vessel; DA: dorsal aorta. Scale bar: 20 μm.

**<u>Video 4: Ablated ISV ECs rapidly activate Erk-signalling following vessel wounding.</u>** Time-lapse video of the ablated ISV in a 4 dpf *Tg(fli1aep:EKC);Tg(fli1a:H2B-mCherry)* larva before (pre-ablation) and after (post-ablation) vessel wounding at indicated timepoints. Post-ablation video starts at 2 minutes post-ablation due to the time taken to transfer the larvae between microscopes and for preparation of imaging. Left panel shows the *fli1aep:EKC* expression and the right panel shows the nuclear *fli1aep:EKC* expression with intensity difference represented in 16 colour LUT (Fiji). The *fli1a:H2B-mCherry* signal was used to mask the nucleus. Z stacks were acquired every 1 minutes for 20 minutes before and after vessel wounding using an Andor Dragonfly Spinning Disc Confocal microscope. Photobleaching was minimised using the bleach correction tool (correction method: Histogram Matching) in FIJI.

ISV: intersegmental vessel. Scale bar: 20 μm.

**<u>Video 5: Adjacent ISV ECs rapidly activate Erk-signalling following vessel wounding.</u>** Time-lapse video of the adjacent ISV in a 4 dpf *Tg(fli1aep:EKC);Tg(fli1a:H2B-mCherry)* larva before (pre-ablation) and after (post-ablation) vessel wounding at indicated timepoints. Post-ablation video starts at 2 minutes post-ablation due to the time taken to transfer the larvae between microscopes and for preparation of imaging. Left panel shows the *fli1aep:EKC* expression and the right panel shows the nuclear *fli1aep:EKC* expression with intensity difference represented in 16 colour LUT (Fiji). The *fli1a:H2B-mCherry* signal was used to mask the nucleus. Z stacks were acquired every 1 minutes for 20 minutes before and after vessel wounding using an Andor Dragonfly Spinning Disc Confocal microscope. Photobleaching was minimised using the bleach correction tool (correction method: Histogram Matching) in FIJI.

ISV: intersegmental vessel. Scale bar: 20 μm.

**<u>Video 6: Skin epithelial and muscle cells do not maintain high Erk activity for 3 hours following muscle wounding.</u>** Time-lapse video of the trunk in a 30 hpf *Tg(ubb:Mmu.Elk1-KTR-mCherry)* embryo. Z stacks were acquired every 21 minutes from 5 mpa until 3 hpa using a Leica SP8 X WLL confocal microscope (n=6 embryos).

Scale bar: 20 μm.

**<u>Video 7: ISVs in 4 dpf larvae do not have active Ca^2+^ signalling.</u>** Time-lapse video of ISVs in a 4 dpf *Tg(actb2:GCaMP6f);Tg(kdrl:mCherry-CAAX)* larva. Left panel shows both the *actb2:GCaMP6f* and the *kdrl:mCherry-CAAX* expressions and the right panel shows the *actb2:GCaMP6f* expression. Z stacks were acquired every minute for 15 minutes using a Leica SP8 confocal microscope.

ISV: intersegmental vessel. Scale bar: 50 μm.

**<u>Video 8: ISVs rapidly activate Ca^2+^ signalling following vessel wounding.</u>** Time-lapse video of both ablated and adjacent ISVs in a 4 dpf *Tg(actb2:GCaMP6f);Tg(kdrl:mCherry-CAAX)* larva following vessel wounding. Left panel shows both the *actb2:GCaMP6f* and the *kdrl:mCherry-CAAX* expressions and the right panel shows the *actb2:GCaMP6f* expression. Z stacks were acquired every minute from 5 mpa until 20 mpa using a Leica SP8 confocal microscope.

ISV: intersegmental vessel. Scale bar: 50 μm.

## Notes

### Competing Interest Statement

The authors have declared no competing interest.

